# Engineered retrons generate genome-independent protein-binding DNA for cellular control

**DOI:** 10.1101/2023.09.27.556556

**Authors:** Geonhu Lee, Jongmin Kim

**Author notes:** Corresponding author. (G.L.); (J.K.).

## Abstract

DNA-protein interactions are core components of myriad natural and synthetic gene networks. Despite the potential new design space, DNA-protein interactions remain underexploited in vivo due to challenges in controlling specific DNA segments, including protein-binding sequences, independently of the genome. Here we engineer retrons, prokaryotic retroelements, to intracellularly generate genome-independent programmable small DNA for sequence-specific protein-binding. Using reprogrammed retron-derived DNA for allosteric transcription factor, we demonstrated dynamic regulation of synthetic gene networks and construction of automated feedback circuits for signal amplification, adaptation, and memory. Furthermore, we developed a new class of stimuli-responsive molecular “bait and prey” that enable modular, rapid, and post-translational control of protein subcellular localization. This work substantially expands possible application area of DNA-protein interactions, laying the foundation for technical advances in synthetic biology.

**One-Sentence Summary:** We demonstrate new ways to control, design, and exploit DNA-protein interactions in living cells using engineered retron-generated small DNA.

## Introduction

Programmable biomolecular interactions between DNA, RNA, and proteins are the main building blocks for synthetic biological systems. Specifically, DNA-protein^1-6^, RNA-protein^7,8^, RNA-RNA^8-10^, and protein-protein^3,11-14^ interactions have been extensively engineered and used to construct genetically encodable synthetic biological devices for a myriad of applications, such as gene regulation and editing^1,3,7^, biosensors^2,4,6^, self-assembled nanostructure^8,11^, complex logic computation^9^, and metabolic engineering^5,8,10^. These intermolecular interactions can be dynamically controlled by modulating the copy-number or state of participating molecules, primarily through gene expression regulation or molecular structural change of RNA or protein components. However, similar approaches for DNA in living cells are challenging even in relatively simple bacteria. In contrast to the proteome and transcriptome, where small subunits are physically separated, subunits of the genome are covalently concatenated within larger DNA molecules, such as chromosomes or plasmids. This restricts the independent control of specific functional parts of DNA, such as short (∼20 bp) protein-binding sequence. Despite DNA-protein interactions have served critical roles since the dawn of synthetic biology with their reliable modularity, tunability, orthogonality, and signal-responsibility, their usage in vivo is still limited to the molecular events placed on the genome. Whereas, in vitro conditions free from these restrictions have allowed wider applications, including comprehensive characterization of DNA-protein interactions ^18^, real-time biosensor^19^, and DNA-protein hybrid nanostructures^20^. Pioneering work by J. Elbaz *et al*. demonstrated unlocking the cellular DNA restrictions by engineering eukaryotic retroviral reverse transcription systems to intracellularly produce DNA oligonucleotides with programmable sequences^21^. However, this system requires relatively heavy genetic components and building complex DNA nanostructures to address low stability and functionality.

Retrons, widespread prokaryotic simple reverse transcription systems composed of a non-coding template RNA and a reverse transcriptase (RT) pair (Fig. 1A) ^22-26^, recently have gained attention due to their compact genetic size, simple mechanism, programmability, and high productivity for single-stranded DNA up to 50,000 copies per cell^27^. Owing to these engineering-tractable properties, retrons have been re-purposed for in situ generation of template DNA for homologous recombination^17,28,29^ and CRISPR-Cas adaptation^30^, advancing dynamic genome engineering in bacteria, yeast^27,28^, and even human cells^29^. Retron-derived DNA (retron-DNA) also has an underexplored characteristic feature, double-stranded hairpin structures commonly found in most retrons discovered so far. This domain lacks evolutionary conserved sequences^31^, suggesting that the retron-DNA can be reprogrammed to mediate sequence-specific protein binding.

**Fig 1.**
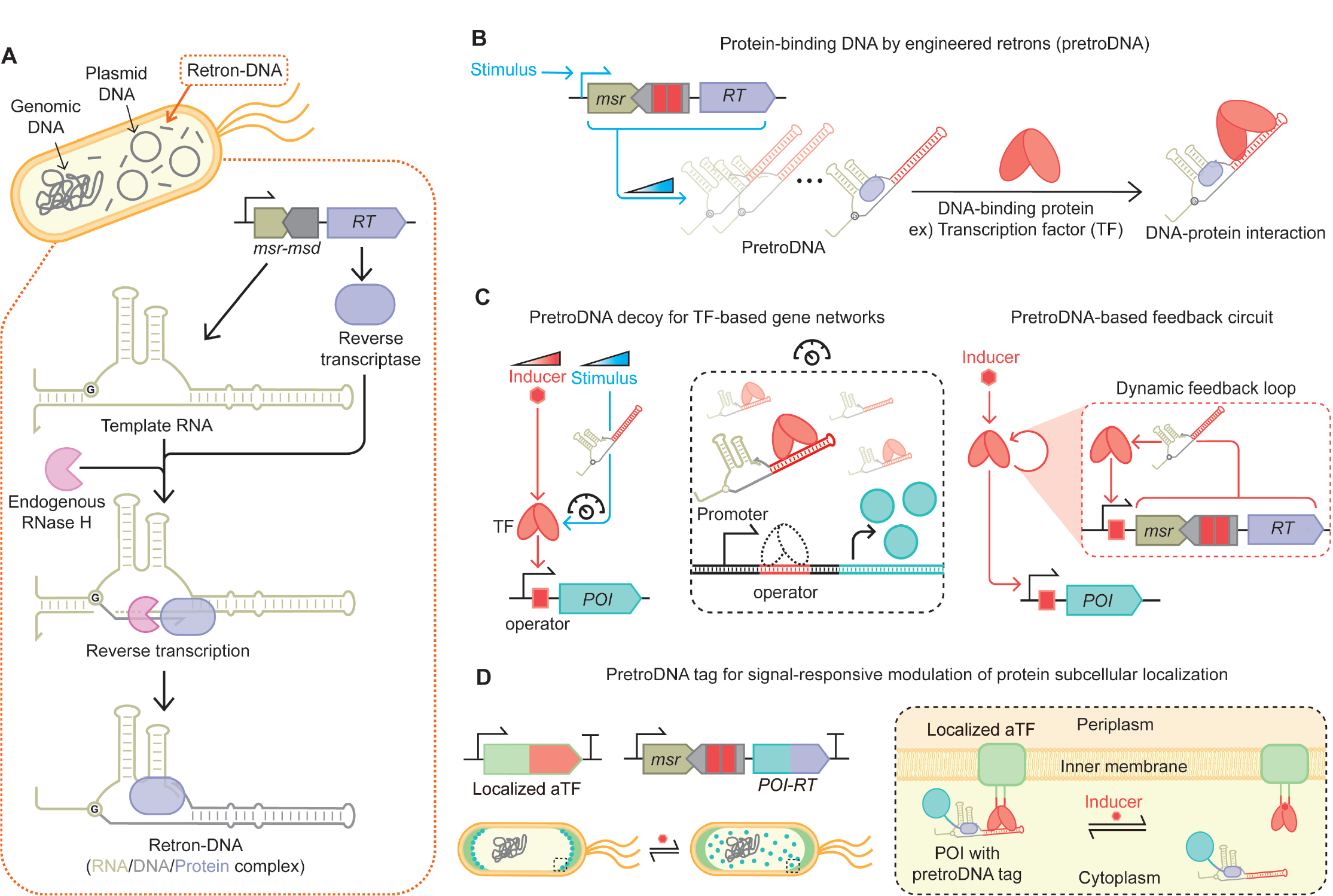
PretroDNA, protein-binding small DNA in situ generated by engineered retrons. (**A**) Bacterial retrons are composed of *msr-msd* and *RT* genes. *Msr-msd* is transcribed into a noncoding template RNA with characteristic structures. The long hairpin domain of template RNA, encoded in *msd*, is converted to retron-DNA by reverse transcriptase and RNase H. The final product is a DNA/RNA/protein complex, in which the noncoding RNA is covalently linked to retron-DNA via a 2’,5’-phosphodiester linkage, and RT remains bound to retron-DNA even after reverse transcription is complete. (**B**) Engineered retrons with reprogrammed *msd* can generate pretroDNA encoding specific protein-binding sequences in their fully double-stranded hairpin domain. The copy-number of intracellular pretroDNA can adjusted by regulating its expression using stimulus-inducible promoters. (**C**) The activity of TF-based gene networks can be dynamically fine-tuned by regulating the expression of pretroDNA encoding TF-binding “operator” sequences. These pretroDNA decoys occupy DNA-binding domain of the TF, thus prevent the TF from binding to the operator sites in the promoter region. Rewiring pretroDNA expression with the target TF-responsive promoter constitutes automated feedback loops that exhibit non-canonical dynamic responses. (**D**) Allosteric TFs (aTFs) sense specific signals and undergo conformational changes that lead to association/dissociation transition with their partner DNA. aTF/pretroDNA pair is a new class of signal-responsive molecular “bait and prey”. Dynamic, rapid, and post-translational modulation of protein subcellular localization is achieved by gene fusion of aTF and RT to an arbitrary localized protein and protein-of-interest (POI), respectively.

Here we develop “Protein-binding small DNA in situ generated by engineered RETROns”, termed pretroDNA (Fig. 1B). This was achieved by reprogramming the double-stranded domain of retron-DNA with specific protein-binding sequences. Using pretroDNA, we demonstrate new ways to control, design, and exploit DNA-protein interaction networks in living cells (Fig. 1 C and D).

## Results

### Programmability of the retron-DNA double-stranded domain

To repurpose retron-DNA for sequence-specific DNA-protein interactions, the stem region should be programmable. To verify this, we used a plasmid encoding the reconstituted Eco1-retron from *Escherichia coli* BL21^17,23^, with the template RNA under IPTG-inducible P_*tac*_ and constitutively expressed RT (Fig. 2A). First, to determine the relative copy-number ratio of retron-DNA to the plasmid, quantitative PCR analysis was conducted with two primer sets, the first anneals to both retron-DNA and *msd* region of the plasmid, while the second anneals only to the plasmid. The ratio of retron to plasmid DNA in *E. coli* with IPTG induction reached 538 (± 31.8 SD) (Fig. 2B), which is 84-fold higher than without IPTG (6.4 ± 1.7 SD), indicating the dynamic copy-number tunability of retron-DNA. Given that the plasmid copy-number with CloDF13 origin of replication is known to be 20∼40 per cell^32^, the retron-DNA would reach 10,760∼21,520 copies/cell.

**Fig 2.**
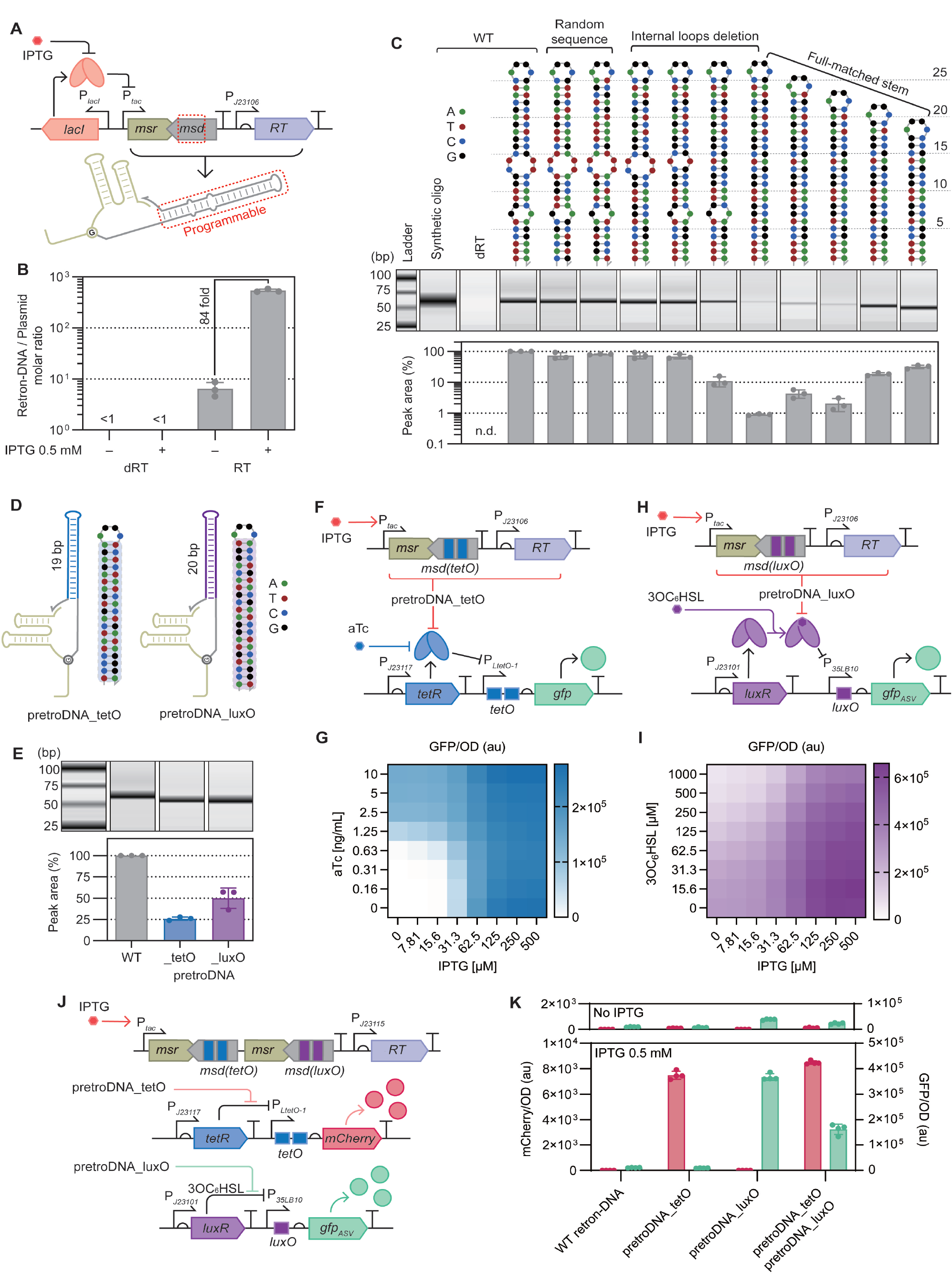
Reprogramming the retron-DNA hairpin domain generates TF-binding pretroDNA. (**A**) Description of pRetro plasmid encoding an IPTG-inducible retron-DNA producing gene circuit. Red boxes indicate the hairpin region to be reprogrammed. (**B**) Quantitative PCR analysis shows that the copy-number of retron-DNA is increased by IPTG in *E. coli* carrying pRetro. (**C**) Capillary electrophoresis analysis of retron-DNA variants produced in *E. coli* deciphers programmability of the hairpin region. The predicted DNA secondary structures-with sequences placed at the top represents the hairpin region of each variant. Peak area represents the relative intracellular productivities compared to WT retron-DNA. Synthetic oligo is chemically synthesized WT retron-DNA. 0.5 mM IPTG was used. n.d., not detected. (**D**) Description of pretroDNA_tetO and pretroDNA_luxO encoding TetR- and LuxR-binding sequences in the hairpin region, respectively. (**E**) Capillary electrophoresis analysis of the pretroDNAs produced in *E. coli*. 0.5 mM IPTG was used. (**F** and **G**) Dynamic fine-tuning of the aTc-responsive TetR/P_*LtetO-1*_ gene circuit by pretroDNA_tetO-mediated TetR decoying. (**H** and **I**) Dynamic fine-tuning of the 3OC_6_HSL-responsive LuxR/P_*35LB10*_ gene circuit by pretroDNA_luxO-mediated LuxR decoying. *ASV* is a degradation tag for quickly synchronizing the tuned transcriptional activity of P_*35LB10*_ with the measured GFP level. (G and I) Values are the mean of n=3 biological replicates. (**J** and **K**) Tandem co-expression of pretroDNA_tetO and pretroDNA_luxO shows the parallel TF-decoying functionality without crosstalk. 1 μM 3OC_6_HSL was used. Error bars indicate SD for n≥3 biological replicates. DNA secondary structure predictions were conducted using NUPACK software^16^. dRT, deactivated RT^17^.

Next, to examine the stem programmability, we generated retron-DNA variants and assessed their productivity (Fig. 2C). Sequence randomizing without structural change resulted in similar productivity compared to WT retron-DNA, whereas interchanging internal loops into hybridized base pairs led to productivity reduction. These results suggest that the copy-number of retron-DNA is mainly determined by structural factors rather than the sequence itself. Since most protein-binding DNAs are fully double-stranded, we focused on the full-matched stem length range over which retron-DNA can be stably produced. Assessment for varied length (17∼25 bp) showed that a severe loss of productivity, 90% reduction compared to WT retron-DNA, occurred at 21 bp or more. The 90% reduced copy-number would be 1,076∼2,125 per cell, which is still higher than the average protein copy-number in *E. coli* (∼1,000 per cell) ^33^. Also, considering most transcription factor (TF)-binding sequences are in the range of 5∼20 bp^34^, the available stem length range would be sufficient. Finally, we confirmed that these engineered retron-DNAs have the designed sequences by analyzing the two stem-randomized variants (fig. S1). Overall, these results demonstrate that the double-stranded domain of retron-DNA has sufficient programmability to be re-encoded with specific protein-binding sequences.

### TF-binding pretroDNA

We then aimed to develop pretroDNA for a representative class of DNA-binding proteins, TFs. Generally, TFs bind to DNA of specific “operator” sequences, usually located within promoter regions, thereby altering the transcriptional activity. TFs bind to several operator sites throughout the genome, and these distributed sites compete for the common TF^35^. Also, distantly inserted operators can sequester the TFs from their cognate promoters, thereby titrating transcriptional activity. Synthetic and endogenous TF-based gene networks have been tuned using these DNA decoys^35-38^. However, dynamically adjusting the copy-number of decoys is challenging, often requiring repetitive cloning to construct a library with varied copy-numbers and subsequent screening for optimization. In contrast, pretroDNA encoding TF-binding sequences could serve as dynamically copy-number tunable DNA decoys (Fig. 1B).

To examine the concept, we chose two target TFs, TetR and LuxR, used in plasmid-based DNA decoy studies ^37^. TetR binds to *tetO* in the P_*LtetO-1*_, thereby repressing downstream gene in the absence of its allosteric inducer, anhydrotetracycline (aTc). In contrast, LuxR is normally detached from *luxO* within P_*35LB10*_^39^, 3-oxohexanoyl-homoserine lactone (3OC_6_HSL) causes LuxR to bind to *luxO* and suppress downstream gene.

First, we generated two pretroDNAs encoding *tetO* (19 bp) or *luxO* (20 bp) sequences (Fig. 2D) and confirmed their intracellular production (Fig. 2E) and sequences (fig. S2). To validate the decoying capacity for TetR, pretroDNA_tetO was co-implemented with a TetR/P_*LtetO-1*_ circuit in which GFP under P_*LtetO-1*_ is repressed by constitutively expressed TetR (Fig. 2 F and G, fig. S3). IPTG induction of pretroDNA_tetO resulted in GFP expression, while basal level of GFP was observed without IPTG. Rest controls showed basal levels of GFP with and without IPTG, confirming the specific decoying capacity (fig. S3, B and C). We then examined the dynamic fine-tuning of the circuit using pretroDNA_tetO, simply varying IPTG concentration generated diverse profiles of aTc-GFP dose-response (Fig. 2G). This combined circuit functions as an OR logic gate that processes two inputs, aTc and pretroDNA_tetO. Finally, we conducted time-course observation of GFP and found that the effect of pretroDNA_tetO was observed approximately 40 minutes after the addition of IPTG (fig. S4).

We also investigated pretroDNA_luxO-mediated LuxR decoying using the similar experimental framework. In the LuxR/P_*35LB10*_ circuit, GFP under P_*35LB10*_ is repressed by 3OC_6_HSL-bound LuxR (Fig. 2H, fig. S5). First, we confirmed the specific decoying capacity of pretroDNA_luxO for LuxR (fig. S5, B and C). Dynamic fine-tuning of the circuit was also achieved by varying IPTG concentration, and this combined circuit functions as a two-input IMPLY gate (Fig. 2I).

Finally, we examined the compatibility and orthogonality of the two pretroDNA using the TetR/P_*LtetO-1*_ circuit for mCherry output and the LuxR/P_*35LB10*_ circuit for GFP output (Fig. 2J). Only when both pretroDNAs were co-expressed were both outputs turned on, while the expression of either pretroDNA alone led to a single cognate output signal (Fig. 2K). Overall, these results confirm that pretroDNA can specifically interact with target proteins, and TF-binding pretroDNAs can serve as a generalizable post-translational TF regulator for dynamic fine-tuning of the TF-based gene networks.

### Dynamic feedback circuit construction with pretroDNA and TF

Next, we sought to harness the dynamic copy-number tunability of pretroDNA decoys to construct more complex gene networks. For example, automated feedback loops can be constructed by rewiring the pretroDNA decoy to the target TF-responsive promoter (Fig. 1C). First, we designed a positive feedback loop in which pretroDNA_tetO under P_*LtetO-1*_ is induced by aTc and subsequently amplifies its own expression by decoying TetR (Fig. 3A). Positive feedback is useful for enhancing sensitivity with controlled leakage in biosensor development^40,41^, so we integrated the pretroDNA_tetO-based feedback with an optimized TetR-based aTc biosensor used in figure 2F and G^37^. This combined biosensor exhibited an order of magnitude improved sensitivity to ∼0.14 ng/ml compared to the dRT control, with negligible background noise (Fig. 3B). We also attempted to tune the feedback strength by engineering the a1/a2 region of template RNA, which is known to affect the retron-DNA productivity^29^. Stronger feedback with an extended a1/a2 (23 bp) resulted in further improved sensitivity to ∼0.05 ng/ml but also increased background noise. These results confirm that pretroDNA-based positive feedback can improve sensitivity of TF-based biosensor and further optimization can be achieved through fine-tuning of the feedback strength.

**Fig 3.**
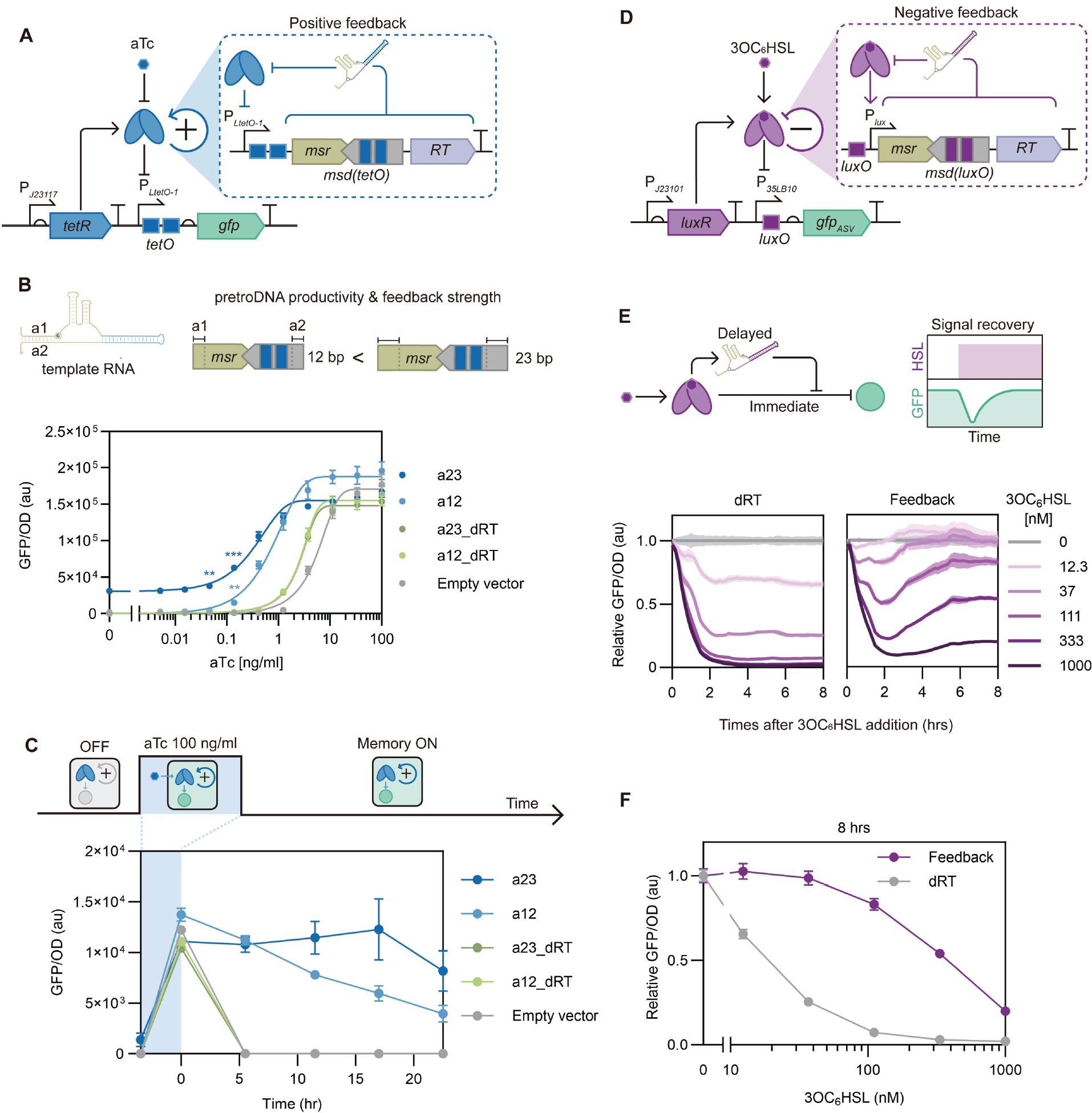
Synthetic automated feedback circuits by rewiring pretroDNA decoy with the target TF-responsive promoter. (**A**) Description of the pretroDNA_tetO-based positive feedback and integration with the TetR/P_*LtetO-1*_-based aTc biosensor circuit. (**B**) The positive feedback improves sensitivity and amplifies the output of aTc biosensor. Extension of the a1/a2 region of the template RNA increases pretroDNA production and strengthens the feedback. **P<0.01 and ***P<0.001 from two-tailed student t-test with the values in no aTc. (**C**) The positive feedback maintains the activation of P_*LtetO-1*_ after the aTc removal, by which transient exposure to aTc is converted into long-term GFP expression. Cells were cultured with aTc for 3.5 hours, then washed with aTc-free medium. Every measurement followed by 1:100 dilution with fresh medium. (**D**) Description of the pretroDNA_luxO-based negative feedback and integration with the LuxR/P_*35LB10*_ circuit. (**E, F**) The negative feedback endows signal recovery response dynamics to the LuxR/P_*35LB10*_ circuit. Error bars indicate SD for n≥3 biological replicates.

Positive feedback is also the foundation of biological memory, extending the lifetime of activated gene networks even after the input signal disappeared^3,42^. Thus, we examined whether pretroDNA_tetO-based positive feedback can convert transient aTc exposure into long-term memory. To test this, the cells were initially cultured with 100 ng/ml aTc for 3.5 hours, then washed and regularly diluted with aTc-free fresh medium. The dRT and empty vector controls showed a rapid decay of GFP to basal level at 5 hours after the aTc removal (Fig. 3C). Whereas, a positive feedback circuit (a12) showed a much slower decay, maintaining 82% and 29% at 5 and 22 hours, respectively. Another circuit with a stronger feedback (a23) showed an extended memory, with no apparent reduction in GFP up to 18 hours.

Encouraged by the achievement of positive feedback, we then constructed a negative feedback loop using pretroDNA_luxO and LuxR and integrate it with the LuxR/P_*35LB10*_ circuit (Fig. 3D). In this combined circuit, 3OC_6_HSL represses GFP under *P*_*35LB10*_ and simultaneously induces pretroDNA_luxO under *P*_*lux*_, a promoter activated by 3OC_6_HSL-bound LuxR. We expected this circuit to show dynamic “decrease-then-recover” responses in GFP expression (Fig. 3E), due to the time differences between the immediate repression of GFP and the delayed pretroDNA_luxO-mediated de-repression. The expected response dynamics were observed in cells with the negative feedback, while dRT control showed only the decrease without recovery (Fig. 3 E and F). Altogether, these results demonstrate that the dynamic copy-number tunability of pretroDNA can be exploited to construct automated feedback circuits by combining with regulatory transcription systems.

### Allosteric TF and pretroDNA pair for signal-responsive modulation of protein subcellular localization

Association/dissociation between allosteric TF (aTF) and the operator DNA can be switched by specific allosteric triggering stimuli, such as biomarkers^6,19^, heavy metals^2^, antibiotics^40^, light^5^, and heat^4^. This universal signal-responsiveness has been primarily exploited for the biosensor development. However, aTF-DNA interactions confined to chromosome or plasmid in bacteria can limit their application scope. In contrast, other signal-responsive biomolecular binding pairs, including protein-protein interactions, have been utilized for a wider range of applications, such as controlling protein subcellular localization^12,13^, despite the narrower pool of available stimulus and orthogonal pairs compared to aTFs-DNA. We speculated that pretroDNA, which is dynamically controllable and physically separated from the genome, can circumvent this limitation.

Thus, we aimed to develop post-translational, dynamic, modular control techniques of protein subcellular localization using aTF/pretroDNA pairs as a new class of signal-responsive molecular “bait and prey”. It was known that RT remains bound to the retron-DNA even after the reverse transcription completed (Fig. 1A) ^25^, suggesting that retron-DNA can be tagged to arbitrary protein “cargo” (Fig 1D). To demonstrate this concept, we fused RT to mCherry, a model cargo, with (GGGGS)_2_ linker and then assessed the RT functionality of the fusion proteins (Fig. 4A). We confirmed the production of pretroDNA_tetO by both N- and C-terminal RT-fused mCherry, indicating RT can tolerate protein fusion at both ends.

**Fig 4.**
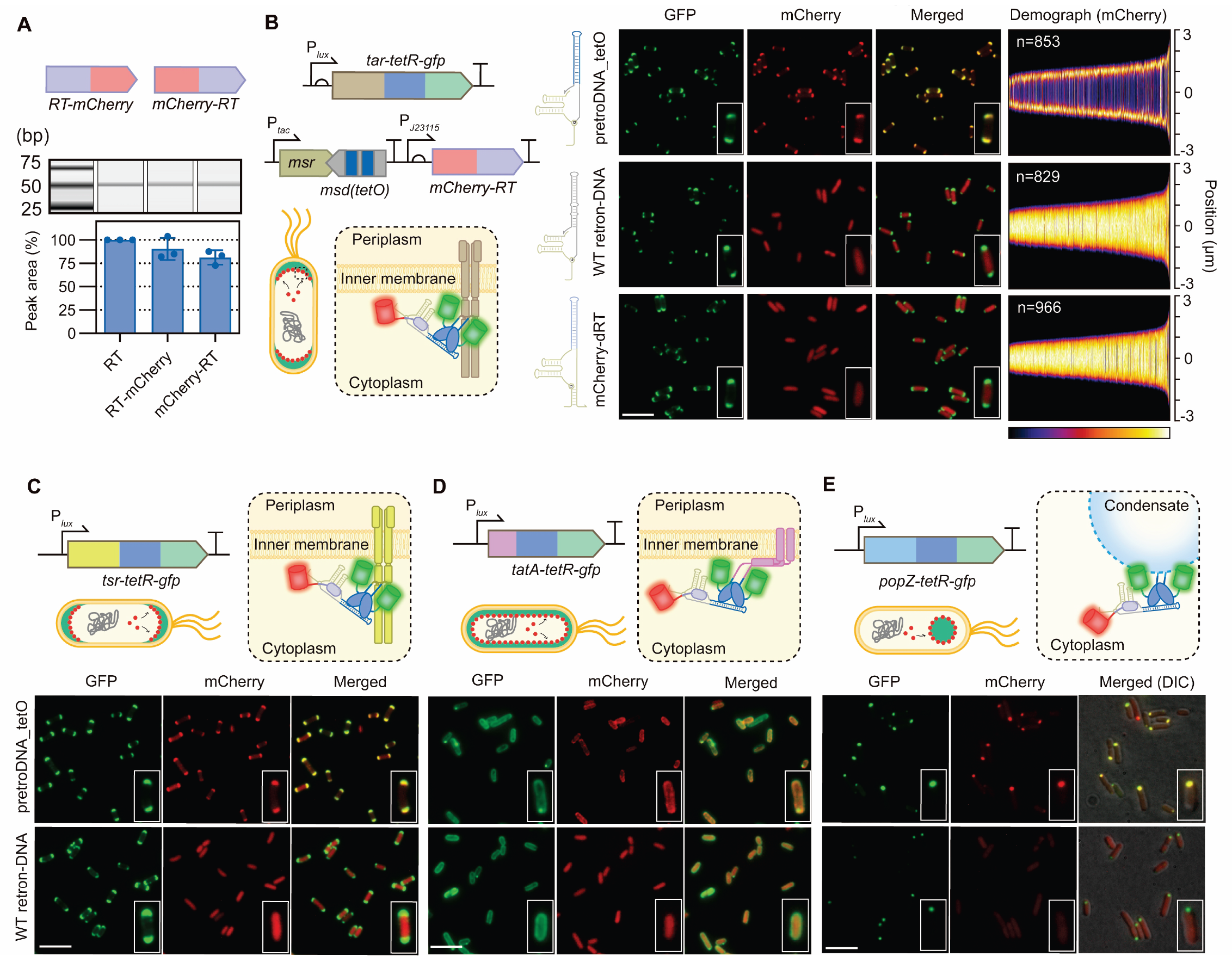
aTF/pretroDNA pair as synthetic molecular “bait and prey” system for modular control of protein subcellular localization. (**A**) PretroDNA_tetO is produced by mCherry-fused RT in *E. coli*. mCherry is a model cargo protein. Error bars indicate SD for n=3 biological replicates. (**B**) mCherry-RT with pretroDNA_tetO tag is recruited to Tar-TetR-GFP localized at membrane poles. Demographs are generated from normalized mCherry intensity profiles along the medial axis of each cell observed in 10 different field of view using MicrobeJ, an ImageJ plugin^15^. Cells are sorted according to their length. (**C** to **E**) mCherry-RT with pretroDNA_tetO tag is co-localized with (**C**) Tsr-TetR-GFP at inner membrane poles, (**D**) TatA-TetR-GFP uniformly distributed throughout the inner membrane, and (**E**) PopZ-TetR-GFP forming condensate at poles. P_*tac*_ was activated by IPTG 0.5 mM. P_*lux*_ was activated by 3OC_6_HSL 2.5 nM (B and C) or 5 nM (D and E). All fluorescence microscopy image data are representatives of n=3 biological replicates. Scale bar, 5 μm. The width of inset in all fluorescence microscopy image data is 2 μm.

Then, we verified whether the mCherry “cargo” could be co-localized with the pre-localized TetR “bait” by the guidance of the pretroDNA_tetO “prey”. As the pre-localized TetR bait, we used Tar-TetR-GFP fusion protein where Tar is an inner membrane-localized chemoreceptor protein concentrated at the poles ^43,44^. Co-localization of GFP and mCherry at the polar membrane was observed in cells expressing pretroDNA_tetO and mCherry-RT (Fig. 4B), while mCherry signals were uniformly distributed throughout the cytoplasm in controls with WT retorn-DNA or mCherry-dRT. These results demonstrate that the localization of mCherry is guided by the specific binding between the TetR domain and pretroDNA_tetO. Similar results were also observed with RT-mCherry, indicating that pretroDNA can be functionally tagged at both ends of a protein (fig. S6). Then, to investigate modularity, we also tested other localizing proteins including Tsr, another chemoreceptor with a similar localization pattern to Tar^43^; TatA, an inner membrane protein known to be evenly distributed throughout the membrane^45^; and PopZ, a condensate-forming protein localized at the poles^46^. Each localizing protein was fused with TetR-GFP and co-expressed with pretroDNA_tetO and RT-mCherry. In all three cases, we observed co-localization of GFP and mCherry with the expected spatial patterns (Fig. 4, C-E), whereas uniformly distributed mCherry signals were observed with WT retron-DNA control.

Finally, we aimed to use the specific stimuli- responsiveness of pretroDNA/aTF pairs for dynamic switching of protein subcellular localization. We chose Tar-TetR-GFP, pretroDNA_tetO, and mCherry-RT to test whether the specific binding between Tar-TetR-GFP and the pretroDNA_tetO/mCherry-RT complex could be reversed post-translationally by aTc (Fig. 5A). To do this, we cultured the cells in LB medium to express all components, then transferred them to M9 minimal medium without carbon sources to attenuate further growth and gene expression prior to the addition of aTc. Fluorescence microscopy observation after 1 hour incubation showed that increasing concentrations of aTc resulted in re-localization of mCherry signal from being concentrated at the poles to a uniform distribution (Fig. 5B). To quantitatively analyze this transition, we calculated the ratio of maximum intensity (*I*_*m*_) to the intensity at the center (*I*_*c*_) from line-scan plot profiles of each cell (Fig. 5A). The *I*_*m*_*/I*_*c*_ ratio would be higher than 1 for concentration at the poles and equal to 1 for a uniform distribution. The ratio gradually decreased from 3.9 (± 2.5, SD) without aTc to 1.1 (± 0.43, SD) with 10 ng/ml aTc (Fig. 5C). Next, to investigate the time required for the re-localization, the cells were observed starting 5 minutes after the addition of aTc (Fig. 5, D and E). Notably, re-localization was almost complete in just 5 minutes, and there was no noticeable change thereafter. This indicates that the transition occurs almost immediately at the post-translational level. Overall, these results demonstrate that the specific stimuli- responsiveness of aTF/pretroDNA pairs can be re-purposed for dynamic modulation of protein subcellular localization.

**Fig 5.**
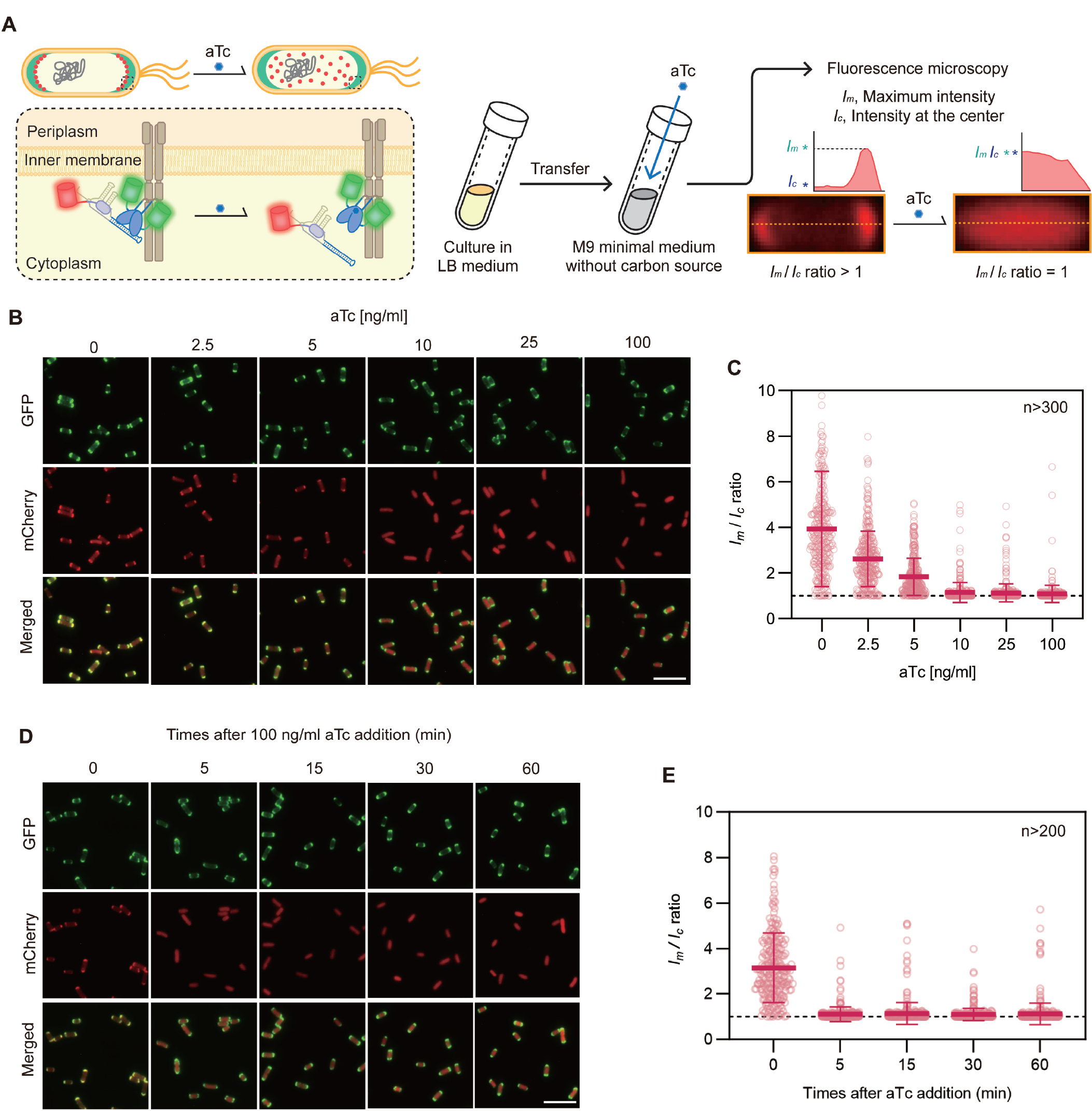
Specific stimuli-responsive dynamic, post-translational modulation of protein subcellular localization using pretroDNA tag. (**A**) Subcellular localization of mCherry-RT with pretroDNA_tetO tag is reversed by aTc from inner membrane poles with Tar-TetR-GFP to a uniform distribution throughout the cells. To analyze the post-translational re-localization, aTc was added to cells in resource-deficient environments followed by fluorescence microscopy observation. *I*_*m*_ and *I*_*c*_ were determined from normalized mCherry intensity profiles along the medial axis of observed cells using MicrobeJ^15^. (**B** and **C**) Fluorescence microscopy (B) and *I*_*m*_/*I*_*c*_ ratio analysis (C) were conducted after 1 hour incubation with aTc. (**D** and **E**) Fluorescence microscopy (D) and *I*_*m*_/*I*_*c*_ ratio analysis (E) by the time of exposure to aTc. All image data are representative of n=3 biological replicates. Scale bar, 5 μm. Error bars indicate SD.

## Discussion

In summary, we found that the long hairpin domain of Eco1 retron-DNA has high programmability, and therefore developed pretroDNA by reprogramming the stem with fully double-stranded, target protein-binding sequences. We demonstrated the expanded capacity for design and control of transcriptional gene network using TF-targeting pretroDNA. Specifically, TetR-and LuxR-based gene circuits were dynamically fine-tuned by simply regulating the cognate pretroDNA decoy expression. Also, by harnessing the dynamic copy-number tunability of pretroDNA, we constructed automated feedback circuits showing dynamic responses, including signal amplification, signal recovery, and sustained activation. This was achieved by simply rewiring pretroDNA expression with the target TF-responsive promoter. Finally, we demonstrated the novel usage of pretroDNA/aTF pair as a new class of synthetic molecular “bait and prey” whose association/dissociation can be dynamically switched by the aTF inducer. By genetically tagging each part to an arbitrary localized protein and the protein-of-interest, we achieved modular, rapid, and post-translational control of protein subcellular localization.

We envision that pretroDNA will expand the overall control capacity and utility of DNA-protein interactions in living cells with three unique properties. First, pretroDNA is a generalizable, effective, and easy-to-use tool that can specifically target and control arbitrary DNA-binding proteins. Second, pretroDNA is physically separated from the genome and therefore independently controllable, expanding the degree-of-freedom for exploiting DNA-protein interactions. Third, pretroDNA can be linked to protein- and potentially RNA-of-interest (Fig. 1A) via gene fusion, allowing synergistic integration with protein- or RNA-based systems.

However, to fully exploit these advantages, several limitations should be addressed. First, generalizability and modularity of pretroDNA should be further verified. We examined only two protein-DNA pairs (TetR/tetO and LuxR/luxO), indicating the need for further testing with other combinations, particularly beyond bacterial aTF. Also, subcellular localization switching of other cargo types, such as enzymes and RNA, should be tested. Second, the detailed relationship between the design of pretroDNA and their copy-number need to be revealed. Although we have demonstrated the importance of structural factors, there is still substantial difference in the copy-number among variants with similar structures. Large scale screening and subsequent establishment of design principles would confer predictability, facilitating development of fit-for-purpose pretroDNA. Third, new design strategies are required to encode longer sequences (>20 bp) for further expanding the application area, such as multiple protein co-localizing scaffolds. This potentially could be achieved through engineering other retron systems ^31^, extending the stem with internal loop insertion, or duplexing the two complementary sequences ^30^.

With these endeavors, pretroDNA would enable wide-ranging novel applications. One direct application area is biosensors. The strategies we demonstrated are generally adoptable to other TF-based biosensors, enabling the development of customized or highly sensitive biosensors for biosensor-assisted laboratory evolution ^47^ or environmental contaminant monitoring ^2,40^. PretroDNA/aTF-mediated localization switching would enable ‘loaded devices’ that are instantly activated upon exposure to specific trigger stimuli ^13^. Their functionality will be determined by their cargo. For example, toxin or recombinase cargo would create kill switch for biocontainment ^48^ or genetic memory for in situ diagnostics ^6^, respectively.

Moreover, integration with sophisticated protein delivery systems, such as engineered bacterial contractile injection ^14^, would enable smart delivery systems that detect the intracellular state of the target and decide whether to release the cargo. Another exciting potential application is DNA-protein hybrid nanoassemblies that can be constructed with modular DNA-binding proteins, such as zinc-finger domains ^20,49^ or CRISPR/Cas systems. These programmable nanostructures can be used as synthetic metabolons or membraneless organelles, and integration of aTF systems would endow the signal-responsiveness. Additionally, retrons function across kingdoms of life, therefore pretroDNA would be applicable to organisms beyond bacteria.

Finally, emerging cellular DNA dynamic control techniques, including retrons and copy-number tunable plasmids ^50^, are unlocking the limited use of DNA in living cells and opening new avenues of synthetic biology and cell engineering. With these technologies, DNA is not merely a pre-determined passive “site” where molecular events are driven by RNA and proteins, but also the active “executors”.

## Acknowledgments

We thank R.M. Murray and S. Oh for thoughtful feedback on this manuscript; Kim Lab members, in particular J. Kim, S. Choi, H. Go, H. Kang, C. Kim, and Y. Choi for comments on this manuscript.

## Author contributions

Conceptualization, Methodology, Investigation, Visualization, Funding acquisition, and Writing – original draft, review & editing: G.L. Funding acquisition, Supervision and Writing – review & editing: J.K.

## Data and materials availability

All data are available in the main text or the supplementary materials.

