## Supplementary material for "Engineered retrons generate genome-independent protein-binding DNA for cellular control": Materials & Methods; Supplementary Figures & Tables

Materials and Methods
Figs. S1 to S6
Tables S1 to S4
References

### Materials and Methods

#### Strains and growth conditions

All cloning, retron-DNA biosynthesis, genetic circuit characterization, and fluorescence microscopy observation experiments were conducted using *E. coli* DH5a. The plasmids used in this study are detailed in Supplementary Table 1, and the combinations of plasmids used for each experiment are listed in Supplementary Table 2. Cells were cultivated in LB (Formedium) at 37 °C with the appropriate antibiotics and inducers. The working concentrations of the antibiotics were as follows: 100 µg ml–1 for ampicillin, 50 µg ml–1 for kanamycin, 25 µg ml–1 for chloramphenicol, and 100 µg ml–1 for spectinomycin. All antibiotics were procured from GoldBio.

For the biosynthesis and extraction of retron-DNA (Fig. 2B, C, E; Fig. 4A; Supplementary Fig. 1 and 2), a single colony on an LB plate was inoculated into 3 ml of LB and shaken at 240 rpm overnight. The cultures were then diluted in a 1:100 ratio in 6 ml of LB and incubated for 2 hours, after which 0.5 mM IPTG was added and the cultures were further incubated for 5 hours. The cultures were then pelleted by centrifugation and stored at -20°C for later retron-DNA extraction.

For genetic circuit characterization, qPCR analysis, and microscopy observation, a single colony of transformants on LB plate was inoculated in LB 300 ul in Deep well plate with breathable sealing film and 1000 rpm shaking for overnight. Then the cultures were diluted 1:100 in LB 200 ul in 96 well black plate with clear bottom (SPL) and incubated for 80 min, then appropriate inducers were added and further incubated. Fluorescence and cell density measurements were conducted at 3 hours after the inducer addition (Fig 2G, I, K). Collecting cells for qPCR analysis was conducted at 5 hours (Fig. 2B). Microscopy observations were conducted at 8 hours (Fig. 4B, C), 2 hours (Fig. 4D), 6 hours (Fig. 4E). Analysis of dynamic protein subcellular localization modulation was conducted at 8 hours (Fig. 5B-E).

For the characterization of genetic circuits, qPCR analysis, and fluorescence microscopy observation, a single colony of transformants on an LB plate was inoculated into 300 µl of LB in a deep well plate with a breathable sealing film and shaken at 1000 rpm overnight. The cultures were then diluted in a 1:100 ratio in 200 µl of LB in a 96-well black plate with a clear bottom (SPL) and incubated for 90 minutes. Appropriate inducers were then added and the cultures were further incubated. Fluorescence and cell density measurements were taken 3 hours after the addition of the inducers (Fig. 2G, I, K). Cells for qPCR analysis were collected at 3 hours (Fig. 2B). Microscopy observations were conducted at 8 hours (Fig. 4B, C; fig S6), 6 hours (Fig. 4D), and 2 hours (Fig. 4E). Cells for dynamic modulation of protein subcellular localization (Fig. 5B-E), were collected at 8 hours.

In the characterization of positive feedback for aTc biosensor sensitivity enhancement (Fig. 3B), the overnight seed culture was diluted in a 1:100 ratio with the addition of aTc in 200 µl of LB medium. This culture was then incubated for 3 hours. Subsequently, measurements were taken for both the fluorescence and cell density of the culture. For the memory characterization (Fig. 3C), the overnight seed culture was diluted in a 1:100 ratio with 100 ng/ml of aTc in 200 µl of LB medium. This was then incubated for 3.5 hours, after which the cells were pelleted and washed with fresh LB medium. The cells were then re-diluted in a 1:100 ratio in fresh LB medium and incubated for an additional 5~6 hours. This was followed by another 1:100 dilution and the process was repeated for 4 times. Measurements of GFP level and OD were conducted immediately prior to each dilution.

In the characterization of negative feedback (Fig. 3, E and F), the overnight seed culture was diluted in a 1:100 ratio in 200 µl of LB medium. This was then incubated for 2.5 hours, after which $3OC_6HSL$ was added. The cells were then further incubated for 16 hours. The GFP level and OD were monitored every 10 minutes.

### Plasmid construction

Plasmids were constructed using standard molecular cloning techniques and Gibson Assembly. Genetic parts used in this study are listed in Supplementary Table 3. All primers were purchased from Bionics (Korea).

Eco1 retron sequences from pFF753, $P_{tac}$ sequences (BBa_K864400), PJ23106 (BBa_J23106), and terminator sequences from pXW117Ptet2-gfp (Addgene #160818) were cloned into pCDFDuet-1 to generate pRetro. *dRT* sequences from pFF758 were cloned into pRetro to generate pRetro_dRT. pFF753 (Addgene #61453) and pFF758 (Addgene #61454) were gifts from T. Lu, MIT. To create variants of pRetro including pPretro_tetO and pPretro_luxO, *msd* variant sequences were cloned into pRetro. pRetro_str_1, pRetro_str_2, and pRetro_str_3 were generated by site-directed mutagenesis from pRetro.

pXW117Ptet2-gfp was used as the TetR/$P_{LtetO\text{-}1}$ circuit plasmid pTet_GFP. ASV degradation tag and $P_{35LB10}$ sequences were cloned into pXW117Ptet2-gfp (Addgene #160819) to generate the LuxR/$P_{35LB10}$ plasmid pLux_GFP. pXW117Ptet2-gfp and pXW101Plux2-gfp were gifts from B. Wang, The University of Edinburgh. *mCherry* sequences sequences were cloned into pSC101 plasmid to generate pTet_mCherry. $P_{LtetO\text{-}1}$ sequences from pXW117Ptet2-gfp, pretroDNA_tetO gene sequences from pPretroDNA_tetO, and *RT* sequences from pFF753 were cloned into pCDFDuet-1 to generate pTet_pretroDNA_tetO. *dRT* sequences from pFF758 were cloned into pTet_pretroDNA_tetO to generate pTet_pretroDNA_tetO_dRT.

pretroDNA_tetO sequences PCR amplified with extended a1/a2 region were cloned into pTet_pretroDNA_tetO and pTet_pretroDNA_tetO_dRT to generate pTet_pretroDNA_tetO_a23 and pTet_pretroDNA_tetO_a23_dRT, respectively. $P_{J23115}$ sequences (BBa_J23115) were cloned into pRetro, pPretro_tetO, and pPretro_luxO to generate pRetro_115, pPretro_tetO_115, and pPretro_luxO_115, respectively. pretroDNA_tetO were PCR amplified with substituted a1/a2 region and cloned into pPretro_luxO_115 to generate pPretro_tetO_luxO_J115.

*tar* sequences were PCR amplified from *E. coli* MG1655, *tetR* and *gfp* sequences from pXW117Ptet2-gfp, and *luxR* and $P_{lux}$ sequences from pXW101Plux2-gfp were cloned into pCDFDuet-1 to generate pTar-TetR-GFP. *tsr* and *tatA* sequences from *E. coli* MG1655 and *popZ* sequences from NSSCP (Addgene #183190) were cloned into pTar-TetR-GFP to generate pTsr-TetR-GFP, pTatA-TetR-GFP, and pPopZ-TetR-GFP, respectively. NSSCP was a gift from H. Huang, National Taiwan University. Retron cassettes sequences from pRetro_115 and mCherry sequences were cloned into pACYCDuet-1 plasmid to generate pRetro_mCherry-RT and pRetro_RT-mCherry.

### Retron-DNA extraction and analysis

To compare the relative productivity of retron-DNA variants to that of WT retron-DNA per cell, equal amounts of cells were harvested based on OD600 measurements for each experiment. DNA was extracted from the pelleted cells using a plasmid miniprep kit (Bionics) and then treated with an RNase A/T1 mix (Thermo Fischer) at 37°C for 30 minutes. The extracted retron-DNA was analyzed by capillary electrophoresis using the Qsep-1 (Bioptic) instrument with the Standard Quantitative Cartridge (S2) and Q-analyzer software (Fig. 2C, E; Fig. 4A). A 25/100 bp Mixed DNA Ladder (Bioneer D-1020) was used as a marker. For sequencing analysis, retron-DNA was separated by agarose electrophoresis and gel extraction, followed by PCR extension and amplification using the Seq_1/2/3 primers (Table S3). Specifically, the extracted retron-DNA and Seq_1 were annealed to each other and extended by polymerase reaction, then amplified by PCR using the Seq_2/3 primers. The resulting PCR products were analyzed by Sanger sequencing (Supplementary Fig. 1 and 2).

**Quantitative PCR**

Relative copy-number of retron-DNA to the encoding plasmid in cells was determined by qPCR analysis with two sets of primers. A region outside the msd site exist only in the plasmid was analyzed using primer set 1 (qPCR_plasmid_F/R), and a region within the msd site exist in both the retron-DNA and the plasmid was analyzed using primer set 2 (qPCR_rtDNA_F/R) (Table S3). Difference in cycle threshold (Ct) between the primer set 1 and 2 was taken as ΔCt value. Standard curve of the ΔCt value to the copy-number ratio was generated by conducting qPCR of synoligo_rtDNA and purified pFF758 plasmid mixtures with a range of copy-number ratio. For analysis of copy-number ratio in *E. coli* cells, 5 ul of cell cultures was collected and mixed with 5 ul of diluted water, then incubated at 95C for 5 min. The boiled cultures were 1:100 diluted with water, 1 ul of which was used for 20 ul reaction with 2X qPCR premix (highQu ORA qPCR Green ROX L Mix) and 0.4 uM of each primer. qPCR was conducted using CFX Opus Real-Time PCR Systems (Bio-rad). The copy-number ratio was calculated from measured ΔCt values with the standard curve. Standard curve generation and raw Ct values are described in Data S1.

**Optical density and fluorescence measurements with plate reader**

The levels of GFP, mCherry, and cell density were measured using a plate reader (Agilent BioTek Synergy H1). The excitation and emission wavelengths for GFP were 479/520 nm, and those for mCherry were 587/620 nm. The absorbance level at 600 nm was measured as the optical density to determine the cell density. Each measurement was conducted with a blank LB medium control of equal volume for normalization. In Figure 3B, the well of the empty vector without aTc was used as a blank to distinguish subtle differences. For continuous time-course analysis (Fig. 3C, E), the fluorescence and optical density levels were measured every 10 minutes with shaking at 800 rpm and incubation at 37°C in the plate reader.

**Fluorescence microscopy observation and analysis**

Agarose pads were prepared using TE buffer with 1% agarose and placed on a slide glass. For live cell microscopy, 1 μl of the cell culture was pipetted onto the agarose pad and covered with a cover glass. Microscopy observations were conducted using a Zeiss Axioscope A1 fluorescence microscope equipped with a 100X Plan-Neofluar oil-immersion objective. The GFP signal was detected using a Zeiss filter set 38 (470/40 nm excitation, 495 nm beam splitter, and 525/50 nm emission filter), while the mCherry signal was detected using a Zeiss filter set 20 (546/12 nm excitation filter, 560 nm beam splitter, and 575-640 emission filter). Digital images were captured using an AxioCam HRm camera. For the dynamic modulation of protein subcellular localization (Fig. 5), cultured cells were pelleted and washed twice with M9 minimal medium (Difco M9 Minimal salts, 5x). The cells were then incubated with aTc and subsequently observed using microscopy.

Brightness adjustment, thresholding, and merging of images were performed using Adobe Photoshop. Images in the same figure were processed under identical conditions. Demographs in figure 4B presenting the subcellular distribution of mCherry fluorescence along the medial axis were generated using MicrobeJ (*46*). Normalized intensity profiles were used. At least 10 images from different fields of view (n=3 biological replicates) were analyzed. To facilitate analysis, clustered cells and mis-detected spots (area <0.01) were manually excluded. The detected cells were then sorted in order of increasing length. $I_m$ and $I_c$ values were determined in the half of the normalized mCherry intensity profiles, generated using MicrobeJ, along the axis of each cell. 3 images from n=3 biological replicates were analyzed.

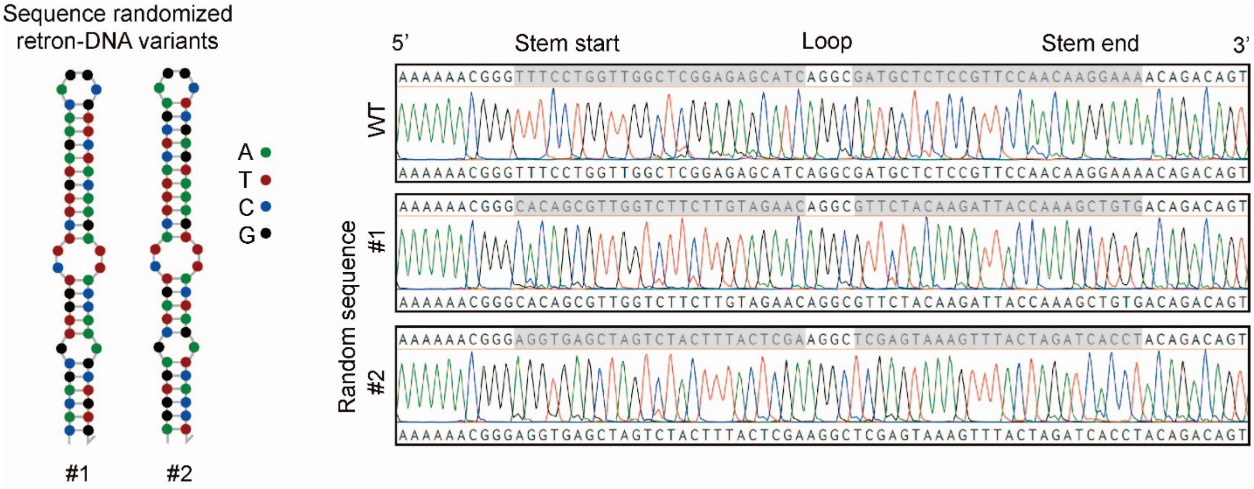


**Fig. S1. Sequence confirmation of stem-randomized retron-DNA variants.** Sanger sequencing analysis results of retron-DNA variants ("Random sequence" in Fig. 2C) produced in *E. coli* are presented. The left side shows the predicted structure of the DNA stem region using NUPACK software, while the right side displays the sequencing results. The top sequences represent the designed sequences, and the bottom sequences represent the analyzed sequences.

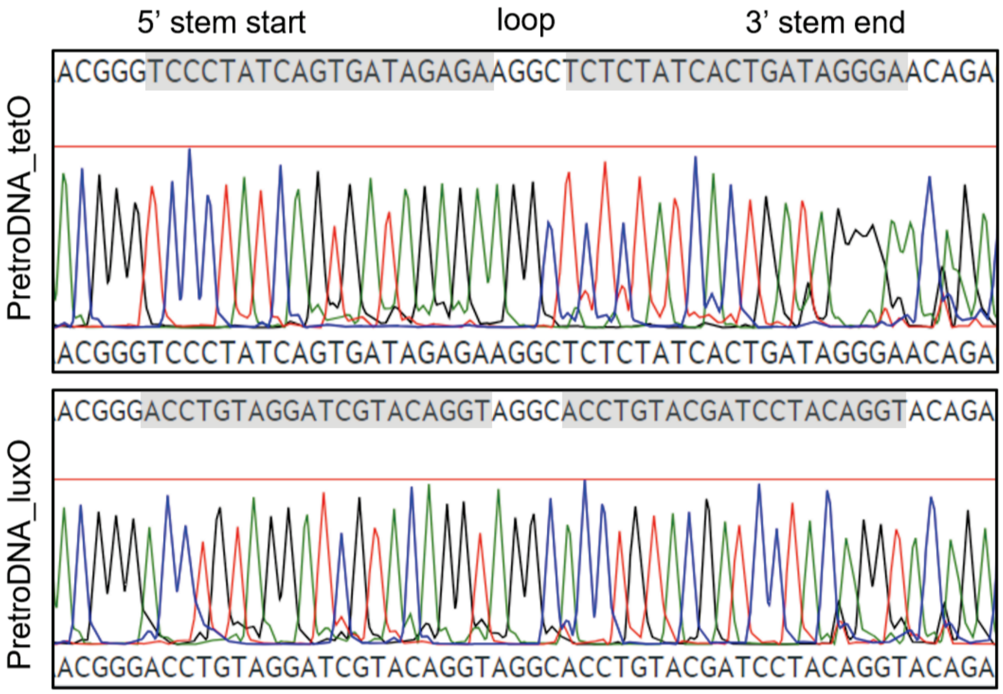


**Fig. S2. Sequence confirmation of pretroDNA_tetO and pretroDNA_luxO.** Sanger sequencing analysis results of pretroDNA produced in *E. coli* are presented. The sequences at the top represent the designed sequences, while the sequences at the bottom represent the analyzed sequences.

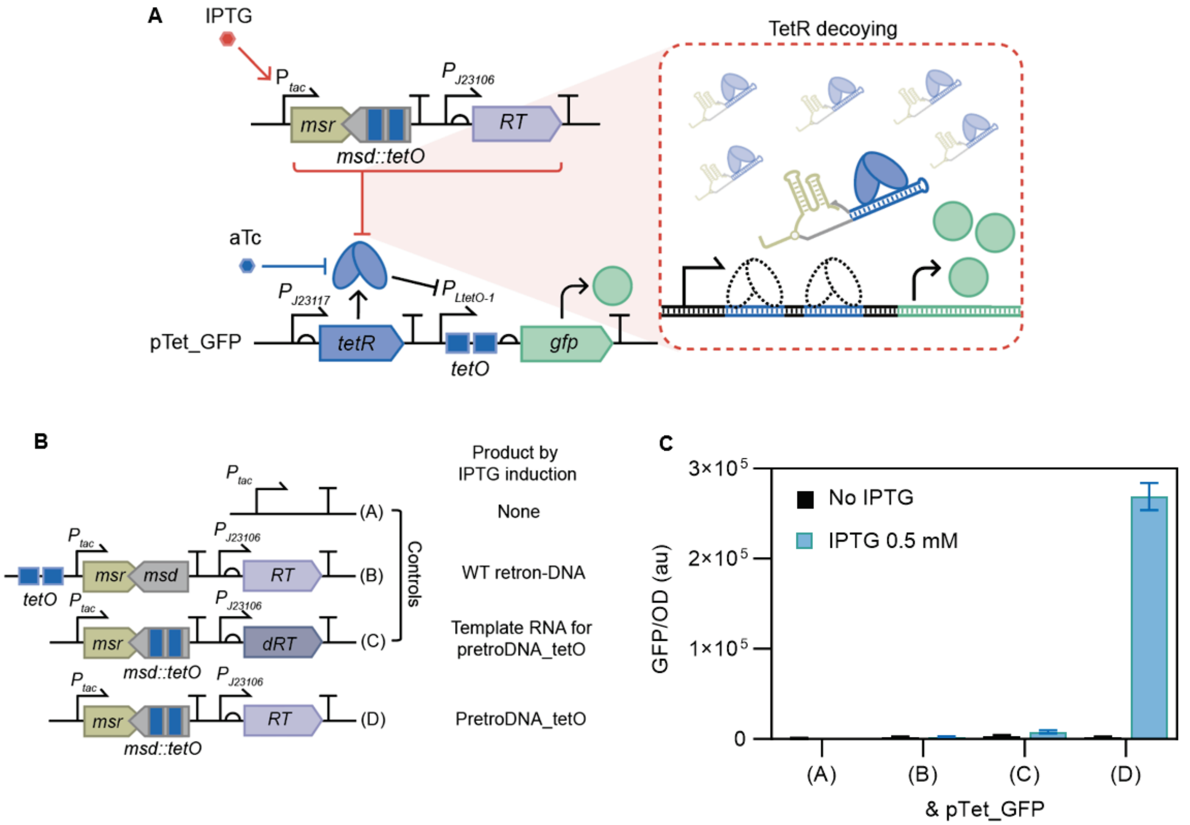


**Fig. S3. PretroDNA_tetO decoys for TetR-regulated gene networks.** (**A**) Scheme depicting the decoying mechanisms of pretroDNA_tetO for TetR. (**B** and **C**) Verification of the specific decoying capacity of pretroDNA_tetO for TetR in *E. coli*. (B) Description of the gene circuit expressing pretroDNA_tetO and control circuits. (C) GFP measurements of cells carrying pTet_GFP and the plasmid described in (B). n=4, biological replicates. Error bars represents SD.

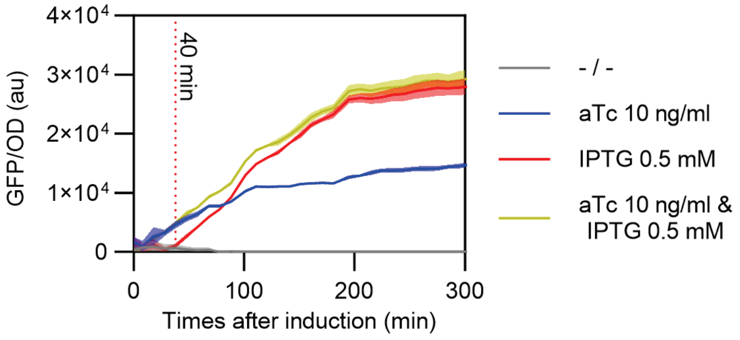


**Fig. S4. Time-course fluorescence measurements of *E. coli* carrying pTet_GFP and pPretro_tetO.** The effect of pretroDNA_tetO emerged at approximately 40 minutes after IPTG induction (Red line). n=4, biological replicates. The filled area represents SD.

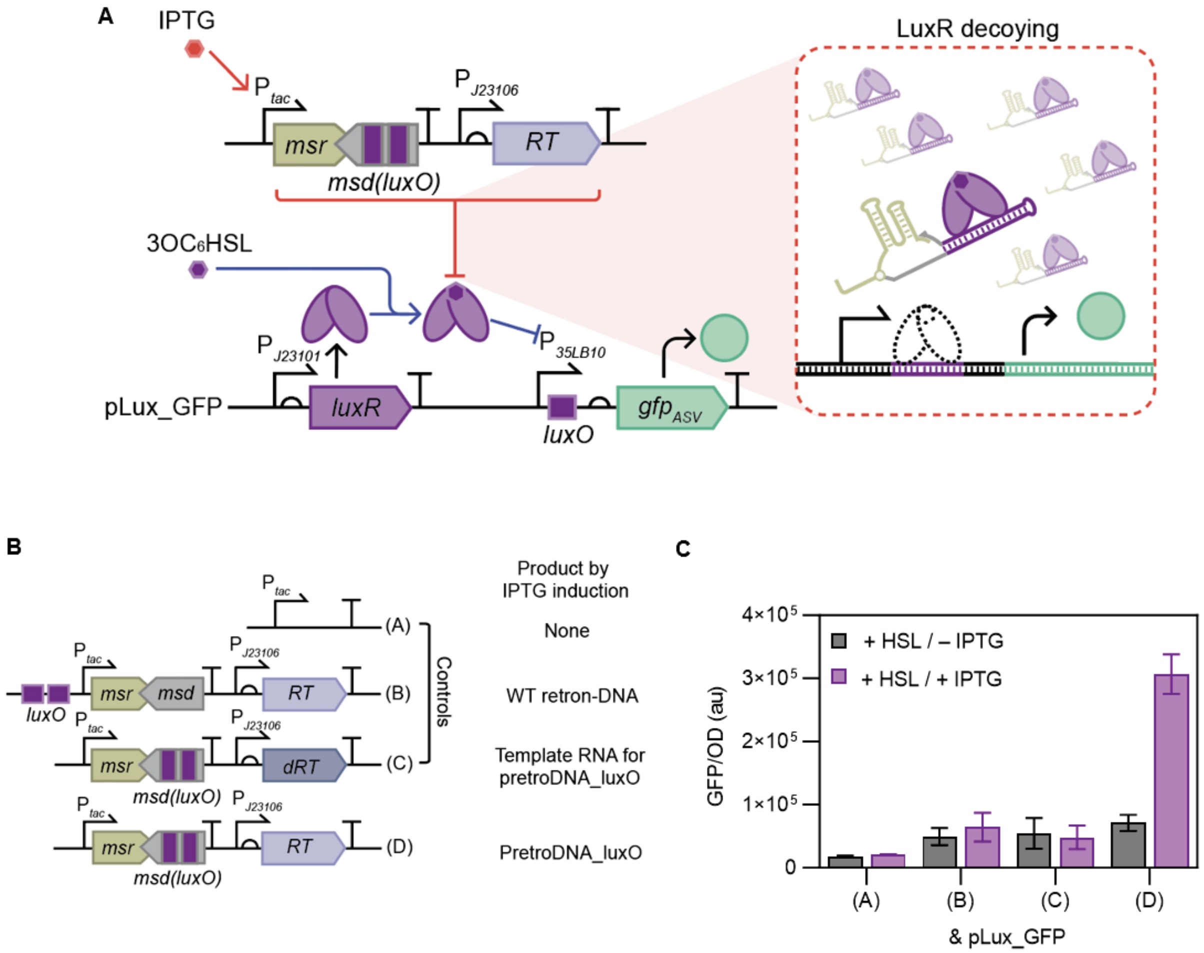


**Fig. S5. PretroDNA_luxO decoys for LuxR-regulated gene networks.** (**A**) Scheme depicting the decoying mechanisms of pretroDNA_luxO for 3OC$_6$HSL-bound LuxR. (**B** and **C**) Verification of the specific decoying capacity of pretroDNA_luxO for 3OC$_6$HSL-bound LuxR in *E. coli*. (**B**) Description of the gene circuit expressing pretroDNA_luxO and control circuits. (C) GFP measurements of cells carrying pTet_GFP and the plasmid in (B). 3OC$_6$SHL, 1 µM. IPTG, 0.5 mM. n=4, biological replicates except for pLux_GFP & plasmid C where n=2 biological replicates due to an experimental error. Error bars represents SD.

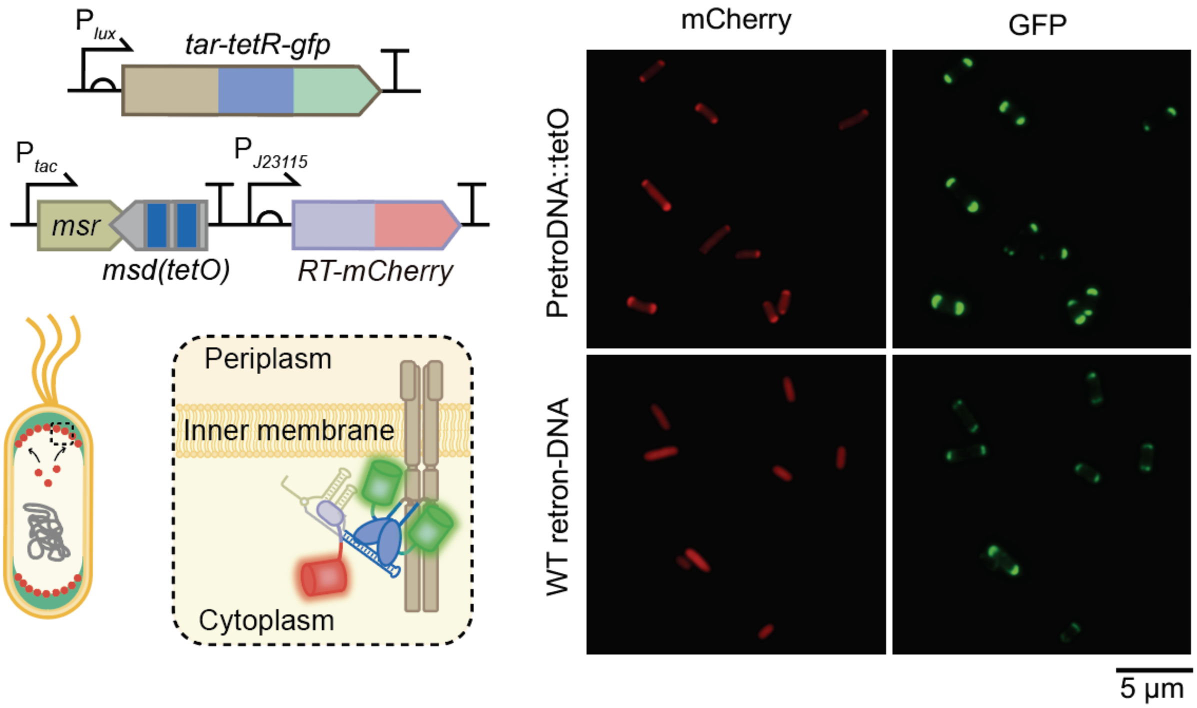


**Fig. S6. RT-mCherry with pretroDNA_tetO tag is recruited to Tar-TetR-GFP localized at membrane poles.** All fluorescence microscopy image data are representatives of n=3 biological replicates. $P_{tac}$ was activated by IPTG 0.5 mM. $P_{lux}$ was activated by $3OC_6HSL$ 2.5 nM.

**Table S1. Plasmids used in this study.**

| Name | Description | Plasmid ori | Antibiotics | Source |
|---|---|---|---|---|
| pFF753 | $P_{lac}$_*msr*_*msd*_*RT* | ColE1 | Chloramphenicol | *(25)* |
| pFF758 | $P_{lac}$_*msr*_*msd*_*dRT* | ColE1 | Chloramphenicol | *(25)* |
| pRetro | *lacI*_$P_{tac}$_*msr*_*msd*_B0015_lambda t0 terminator_$P_{J23106}$_*RT*_T7 terminator | CloDF13 | Spectinomycin | This study |
| pRetro_dRT | *lacI*_$P_{tac}$_*msr*_*msd*_B0015_lambda t0 terminator_$P_{J23106}$_*dRT*_T7 terminator | CloDF13 | Spectinomycin | This study |
| pRetro_17bp | *lacI*_$P_{tac}$_*msr*_*msd(17)*_B0015_lambda t0 terminator_$P_{J23106}$_*RT*_T7 terminator | CloDF13 | Spectinomycin | This study |
| pRetro_19bp | *lacI*_$P_{tac}$_*msr*_*msd(19)*_B0015_lambda t0 terminator_$P_{J23106}$_*RT*_T7 terminator | CloDF13 | Spectinomycin | This study |
| pRetro_21bp | *lacI*_$P_{tac}$_*msr*_*msd(21)*_B0015_lambda t0 terminator_$P_{J23106}$_*RT*_T7 terminator | CloDF13 | Spectinomycin | This study |
| pRetro_23bp | *lacI*_$P_{tac}$_*msr*_*msd(23)*_B0015_lambda t0 terminator_$P_{J23106}$_*RT*_T7 terminator | CloDF13 | Spectinomycin | This study |
| pRetro_25bp | *lacI*_$P_{tac}$_*msr*_*msd(25)*_B0015_lambda t0 terminator_$P_{J23106}$_*RT*_T7 terminator | CloDF13 | Spectinomycin | This study |
| pRetro_rand_1 | *lacI*_$P_{tac}$_*msr*_*msd(rand_1)*_B0015_lambda t0 terminator_$P_{J23106}$_*RT*_T7 terminator | CloDF13 | Spectinomycin | This study |
| pRetro_rand_2 | *lacI*_$P_{tac}$_*msr*_*msd(rand_2)*_B0015_lambda t0 terminator_$P_{J23106}$_*RT*_T7 terminator | CloDF13 | Spectinomycin | This study |
| pRetro_str_1 | *lacI*_$P_{tac}$_*msr*_*msd(str_1)*_B0015_lambda t0 terminator_$P_{J23106}$_*RT*_T7 terminator | CloDF13 | Spectinomycin | This study |
| pRetro_str_2 | *lacI*_$P_{tac}$_*msr*_*msd(str_2)*_B0015_lambda t0 terminator_$P_{J23106}$_*RT*_T7 terminator | CloDF13 | Spectinomycin | This study |
| pRetro_str_3 | *lacI*_$P_{tac}$_*msr*_*msd(str_3)*_B0015_lambda t0 terminator_$P_{J23106}$_*RT*_T7 terminator | CloDF13 | Spectinomycin | This study |
| pEmpty | *lacI*_$P_{tac}$_T7 terminator | CloDF13 | Spectinomycin | This study |
| pPretro_tetO | *lacI*_$P_{tac}$_*msr*_*msd(tetO)*_B0015_lambda t0 terminator_$P_{J23106}$_*RT*_T7 terminator | CloDF13 | Spectinomycin | This study |
| pPretro_tetO_dRT | *lacI*_$P_{tac}$_*msr*_*msd(tetO)*_B0015_lambda t0 terminator_$P_{J23106}$_*dRT*_T7 terminator | CloDF13 | Spectinomycin | This study |
| pRetro_decoy_tetO | *2XtetO*_*lacI*_$P_{tac}$_*msr*_*msd*_B0015_lambda t0 terminator_$P_{J23106}$_*RT*_T7 terminator | CloDF13 | Spectinomycin | This study |
| pPretro_luxO | *lacI*_$P_{tac}$_*msr*_*msd(luxO)*_B0015_lambda t0 terminator_$P_{J23106}$_*RT*_T7 terminator | CloDF13 | Spectinomycin | This study |
| pPretro_luxO_dRT | *lacI*_$P_{tac}$_*msr*_*msd(luxO)*_B0015_lambda t0 terminator_$P_{J23106}$_*dRT*_T7 terminator | CloDF13 | Spectinomycin | This study |
| pRetro_decoy_luxO | *2XluxO*_*lacI*_$P_{tac}$_*msr*_*msd*_B0015_lambda t0 terminator_$P_{J23106}$_*RT*_T7 terminator | CloDF13 | Spectinomycin | This study |
| pXW101Plux2-gfp (pTet_GFP) | $P_{J23117}$_B0030_*tetR*_B0015_$P_{LtetO-1}$_B0030_*gfp*_B0015 | RK2 | Ampicillin | *(35)* |
| pXW101Plux2-gfp | $P_{J23101}$_B0030_*luxR*_B0015_$P_{lux}$_B0030_*gfp*_B0015 | RK2 | Ampicillin | *(35)* |
| pLux_GFP | $P_{J23101}$_B0030_*luxR*_B0015_$P_{35LB10}$_B0030_*gfp*$_{ASV}$_B0015 | RK2 | Ampicillin | This study |
| pTet_mCherry | $P_{J23117}$_B0030_*tetR*_B0015_$P_{LtetO-1}$_B0030_*mCherry*_T7 terminator | SC101 | Kanamycin | This study |
| pRetro_115 | *lacI*_$P_{tac}$_*msr*_*msd*_B0015_lambda t0 terminator_$P_{J23115}$_*RT*_T7 terminator | CloDF13 | Spectinomycin | This study |

| | | | | |
|---|---|---|---|---|
| pPretro_tetO_115 | *lacI*_$P_{tac}$_*msr_msd(tetO)*_B0015_lambda t0 terminator_$P_{J23115}$_*RT*_T7 terminator | CloDF13 | Spectinomycin | This study |
| pPretro_luxO_115 | *lacI*_$P_{tac}$_*msr_msd(luxO)*_B0015_lambda t0 terminator_$P_{J23115}$_*RT*_T7 terminator | CloDF13 | Spectinomycin | This study |
| pPretro_tetO_luxO_115 | *lacI*_$P_{tac}$_*msr_msd(tetO)_msr_msd(luxO)*_B0015_lambda t0 terminator_$P_{J23115}$_*RT*_T7 terminator | CloDF13 | Spectinomycin | This study |
| pTet_pretroDNA_tetO | $P_{LtetO-1}$_*msr_msd(tetO)_RT*_T7 terminator | CloDF13 | Spectinomycin | This study |
| pTet_pretroDNA_tetO_dRT | $P_{LtetO-1}$_*msr_msd(tetO)_dRT*_T7 terminator | CloDF13 | Spectinomycin | This study |
| pTet_pretroDNA_tetO_a23 | $P_{LtetO-1}$_*a1(23bp)_msr_msd(tetO)_a2(23bp)_RT*_T7 terminator | CloDF13 | Spectinomycin | This study |
| pTet_pretroDNA_tetO_a23_dRT | $P_{LtetO-1}$_*a1(23bp)_msr_msd(tetO)_a2(23bp)_dRT*_T7 terminator | CloDF13 | Spectinomycin | This study |
| pLux_pretroDNA_luxO | $P_{lux}$_*msr_msd(luxO)_RT*_T7 terminator | CloDF13 | Spectinomycin | This study |
| pLux_pretroDNA_luxO_dRT | $P_{lux}$_*msr_msd(luxO)_dRT*_T7 terminator | CloDF13 | Spectinomycin | This study |
| pPretro_tetO_106_RT | *lacI*_$P_{tac}$_*msr_msd(tetO)*_B0015_lambda t0 terminator_$P_{J23106}$_*RT*_T7 terminator | p15a | Chloramphenicol | This study |
| pPretro_tetO_106_mCherry-RT | *lacI*_$P_{tac}$_*msr_msd(tetO)*_B0015_lambda t0 terminator_$P_{J23106}$_*mCherry-RT*_T7 terminator | p15a | Chloramphenicol | This study |
| pPretro_tetO_106_RT-mCherry | *lacI*_$P_{tac}$_*msr_msd(tetO)*_B0015_lambda t0 terminator_$P_{J23106}$_*RT-mCherry*_T7 terminator | p15a | Chloramphenicol | This study |
| pPretro_tetO_mCherry-RT | *lacI*_$P_{tac}$_*msr_msd(tetO)*_B0015_lambda t0 terminator_$P_{J23115}$_*mCherry-RT*_T7 terminator | p15a | Chloramphenicol | This study |
| pRetro_mCherry-RT | *lacI*_$P_{tac}$_*msr_msd*_B0015_lambda t0 terminator_$P_{J23115}$_*mCherry-RT*_T7 terminator | p15a | Chloramphenicol | This study |
| pPretro_tetO_mCherry-dRT | *lacI*_$P_{tac}$_*msr_msd(tetO)*_B0015_lambda t0 terminator_$P_{J23115}$_*mCherry-dRT*_T7 terminator | p15a | Chloramphenicol | This study |
| pPretro_tetO_RT-mCherry | *lacI*_$P_{tac}$_*msr_msd(tetO)*_B0015_lambda t0 terminator_$P_{J23115}$_*RT-mCherry*_T7 terminator | p15a | Chloramphenicol | This study |
| pRetro_RT-mCherry | *lacI*_$P_{tac}$_*msr_msd*_B0015_lambda t0 terminator_$P_{J23115}$_*RT-mCherry*_T7 terminator | p15a | Chloramphenicol | This study |
| pTar-TetR-GFP | $P_{J23101}$_B0030_*luxR*_B0015_$P_{lux}$_B0030_*tar-tetR-gfp*_T7 terminator | CloDF13 | Spectinomycin | This study |
| pTsr-TetR-GFP | $P_{J23101}$_B0030_*luxR*_B0015_$P_{lux}$_B0030_*tsr-tetR-gfp*_T7 terminator | CloDF13 | Spectinomycin | This study |
| pTatA-TetR-GFP | $P_{J23101}$_B0030_*luxR*_B0015_$P_{lux}$_B0030_*tatA-tetR-gfp*_T7 terminator | CloDF13 | Spectinomycin | This study |
| pPopZ-TetR-GFP | $P_{J23101}$_B0030_*luxR*_B0015_$P_{lux}$_B0030_*popZ-tetR-gfp*_T7 terminator | CloDF13 | Spectinomycin | This study |
| NSSCP | $J_{23116}$_mRFP-popZ | pMB1 | Chloramphenicol | *(51)* |

**Table S2. Plasmid combinations used in figures.**

| Figure number | Plasmid combination |
|---|---|
| 2B | - pRetro<br>- pRetro_dRT<br>- pFF753<br>- pFF758 |
| 2C | - pRetro_dRT<br>- pRetro<br>- pRetro_rand_1<br>- pRetro_rand_2<br>- pRetro_str_1<br>- pRetro_str_2<br>- pRetro_str_3<br>- pRetro_25bp<br>- pRetro_23bp<br>- pRetro_21bp<br>- pRetro_19bp<br>- pRetro_17bp |
| 2E | - pRetro<br>- pPretro_tetO<br>- pPretro_luxO |
| 2G | - pTet_GFP & pPretro_tetO |
| 2I | - pLux_GFP & pPretro_luxO |
| 2K | - pTet_mCherry & pLux_GFP & pRetro_115<br>- pTet_mCherry & pLux_GFP & pPretro_tetO_115<br>- pTet_mCherry & pLux_GFP & pPretro_luxO_115<br>- pTet_mCherry & pLux_GFP & pPretro_tetO_luxO_115 |
| 3B, C | - pTet_GFP & pEmpty<br>- pTet_GFP & pTet_pretroDNA_tetO_dRT<br>- pTet_GFP & pTet_pretroDNA_tetO_a23_dRT<br>- pTet_GFP & pTet_pretroDNA_tetO<br>- pTet_GFP & pTet_pretroDNA_tetO_a23 |
| 3E, F | - pLux_GFP & pLux_pretroDNA_luxO_dRT<br>- pLux_GFP & pLux_pretroDNA_luxO |
| 4A | - pPretro_tetO_106_RT<br>- pPretro_tetO_106_mCherry-RT<br>- pPretro_tetO_106_RT-mCherry |
| 4B | - pTar-TetR-GFP & pPretro_tetO_mCherry-RT<br>- pTar-TetR-GFP & pRetro_mCherry-RT<br>- pTar-TetR-GFP & pPretro_tetO_mCherry-dRT |
| 4C | - pTsr-TetR-GFP & pPretro_tetO_mCherry-RT<br>- pTsr-TetR-GFP & pRetro_mCherry-RT |
| 4D | - pTatA-TetR-GFP & pPretro_tetO_mCherry-RT<br>- pTatA-TetR-GFP & pRetro_mCherry-RT |
| 4E | - pPopZ-TetR-GFP & pPretro_tetO_mCherry-RT<br>- pPopZ-TetR-GFP & pRetro_mCherry-RT |

| | |
|---|---|
| 5B-E | - pTar-TetR-GFP & pPretro_tetO_mCherry-RT |
| S1 | - pRetro<br>- pRetro_rand_1<br>- pRetro_rand_2 |
| S2 | - pPretro_tetO<br>- pPretro_luxO |
| S3 B, C | - pTet_GFP & pEmpty<br>- pTet_GFP & pPretro_tetO_dRT<br>- pTet_GFP & pRetro_decoy_tetO<br>- pTet_GFP & pPretro_tetO |
| S4 | - pTet_GFP & pPretro_tetO |
| S5 B, C | - pLux_GFP & pEmpty<br>- pLux_GFP & pPretro_luxO_dRT<br>- pLux_GFP & pRetro_decoy_luxO<br>- pLux_GFP & pPretro_luxO |
| S6 | - pTar-TetR-GFP & pPretro_tetO_RT-mCherry<br>- pTar-TetR-GFP & pRetro_RT-mCherry |

### Table S3. Genetic parts sequences.

| Part name | Sequence |
|---|---|
| $P_{J23101}$ | tttacagctagctcagtcctaggtattatgctagc |
| $P_{J23106}$ | tttacggctagctcagtcctaggtatagtgctagc |
| $P_{J23117}$ | ttgacagctagctcagtcctagggattgtgctagc |
| $P_{J23115}$ | tttatagctagctcagcccttggtacaatgctagc |
| $P_{LtetO\text{-}1}$ | tccctatcagtgatagagattgacatccctatcagtgatagagatactgagcacatat |
| $P_{35LB10}$ | ttgacacctgtaggatcgtacaggtataat |
| $P_{tac}$ | gagctgttgacaattaatcatcggctcgtataatgtgtggaattgtgagcggataacaatt |
| $P_{lux}$ | agacctgtaggatcgtacaggtttacgcaagaaaatggtttgttactttcgaataaa |
| B0030 | attaaagaggagaaa |
| B0032 | tcacacaggaaag |
| B0015 | ccaggcatcaaataaaacgaaaggctcagtcgaaagactgggcctttcgttttatctgttgtttgtcggtgaacgctctctactagagtcaca<br>ctggctcaccttcgggtgggcctttctgcgtttata |
| Lambda t0 terminator | gactcctgttgatagatccagtaatgacctcagaactccatctggatttgttcagaacgctcggttgccgccgggcgttttttattggtgagaat |
| T7 terminator | tagcataaccccttggggcctctaaacgggtcttgaggggttttttg |
| *msr* | tgcgcacccttagcgagaggtttatcattaaggtcaacctctggatgttgtttcggcatcctgcattgaatctgagttact |
| *msd* | tctgagttactgtctgttttccttgttggaacggagagcatcgcctgatgctctccgagccaaccaggaaacccgttttttctgac |
| *msd(17)* | tctgagttactgtctgttttcctggttggctcgggcctccgagccaaccaggaaacccgttttttctgac |
| *msd(19)* | tctgagttactgtctgttttcctggttggctcggaggcctctccgagccaaccaggaaacccgttttttctgac |
| *msd(21)* | tctgagttactgtctgttttcctggttggctcggagaggcctctctccgagccaaccaggaaacccgttttttctgac |
| *msd(23)* | tctgagttactgtctgttttcctggttggctcggagagcagccttgctctccgagccaaccaggaaacccgttttttctgac |
| *msd(25)* | tctgagttactgtctgttttcctggttggctcggagagcatcgcctgatgctctccgagccaaccaggaaacccgttttttctgac |
| *msd(str_1)* | tctgagttactgtctgttttcctggttggaacggagagcatcgcctgatgctctccgagccaaccaggaaacccgttttttctgac |
| *msd(str_2)* | tctgagttactgtctgttttccttgttggcacggagagcatcgcctgatgctctccgagccaaccaggaaacccgttttttctgac |
| *msd(str_3)* | tctgagttactgtctgttttccttgttggctcggagagcatcgcctgatgctctccgagccaaccaggaaacccgttttttctgac |
| *msd(rand_1)* | tctgagttactgtctgtcacagctttggtaatcttgtagaacgcctgttctacaagaagaccaacgctgtgcccgttttttctgac |
| *msd(rand_2)* | tctgagttactgtctgtaggtgatctagtaaactttactcgagccttcgagtaaagtagactagctcacctcccgttttttctgac |
| *msd(tetO)* | tctgagttactgtctgttccctatcagtgatagagagccttctctatcactgatagggacccgttttttctgac |
| *msd(luxO)* | tctgagttactgtctgtacctgtaggatcgtacaggtgcctacctgtacgatcctacaggtcccgttttttctgac |
| *a1(23bp)_msr_msd(tetO)_a2 (23bp)* | aagattccgtatgcgcacccttagcgagaggtttatcattaaggtcaacctctggatgttgtttcggcatcctgcattgaatctgagttactgtct<br>gttccctatcagtgatagagagccttctctatcactgatagggacccgttttttctgacgtaagggtgcgcatacggaatctt |
| *RT* | atgaaatccgctgaatatttgaacacttttagattgagaaatctcggcctacctgtcatgaacaatttgcatgacatgtctaaggcgactcgcat<br>atctgttgaaacacttcggttgttaatctatacagctgattttcgctataggatctacactgtagaaaagaaaggcccagagaagagaatgag<br>aaccatttaccaaccttctcgagaacttaaagccttacaaggatgggttctacgtaacattttagataaactgtcgtcatctcctttttctattgga<br>tttgaaaagcaccaatctattttgaataatgctaccccgcatattggggcaaactttatactgaatattgatttggaggatttttttcccaagtttaa<br>ctgctaacaaagtttttggagtgttccattctcttggttataatcgactaatatcttcagttttgacaaaaatatgttgttataaaaatctgctaccac<br>aaggtgctccatcatcacctaaattagctaatctaatatgttctaaacttgattatcgtattcagggttatgcaggtagtcggggcttgatatata<br>cgagatatgccgatgatctcaccttatctgcacagtctatgaaaaaggttgttaaagcacgtgatttttttattttctataatcccaagtgaaggat<br>tggttattaactcaaaaaaaacttgtattagtgggcctcgtagtcagaggaaagttacaggtttagttatttcacaagagaaagttgggatagg<br>tagagaaaaatataaagaaattagagcaaagatacatcatatattttgcggtaagtcttctgagatagaacacgttaggggatggttgtcattt<br>attttaagtgtggattcaaaaagccataggagattaataacttatattagcaaattagaaaaaaaatatggaaagaaccctttaaataaagcga<br>agacctaa |
| *dRT* | atgaaatccgctgaatatttgaacacttttagattgagaaatctcggcctacctgtcatgaacaatttgcatgacatgtctaaggcgactcgcat<br>atctgttgaaacacttcggttgttaatctatacagctgattttcgctataggatctacactgtagaaaagaaaggcccagagaagagaatgag<br>aaccatttaccaaccttctcgagaacttaaagccttacaaggatgggttctacgtaacattttagataaactgtcgtcatctcctttttctattgga<br>tttgaaaagcaccaatctattttgaataatgctaccccgcatattggggcaaactttatactgaatattgatttggaggatttttttcccaagtttaa<br>ctgctaacaaagtttttggagtgttccattctcttggttataatcgactaatatcttcagttttgacaaaaatatgttgttataaaaatctgctaccac<br>aaggtgctccatcatcacctaaattagctaatctaatatgttctaaacttgattatcgtattcagggttatgcaggtagtcggggcttgatatata<br>cgagatatgccgctgctctcaccttatctgcacagtctatgaaaaaggttgttaaagcacgtgatttttttattttctataatcccaagtgaaggat<br>tggttattaactcaaaaaaaacttgtattagtgggcctcgtagtcagaggaaagttacaggtttagttatttcacaagagaaagttgggatagg<br>tagagaaaaatataaagaaattagagcaaagatacatcatatattttgcggtaagtcttctgagatagaacacgttaggggatggttgtcattt |

| | |
|---|---|
| | attttaagtgtggattcaaaaagccataggagattaataacttatattagcaaattagaaaaaaaatatggaaagaaccctttaaataaagcgaagacctaa |
| *tetR* | atgtccagattagataaaagtaaagtgattaacagcgcattagagctgcttaatgaggtcggaatcgaaggtttaacaacccgtaaactcgcccagaagctaggtgtagagcagcctacattgtattggcatgtaaaaaataagcgggctttgctcgacgccttagccattgagatgttagataggcaccatactcacttttgccctttagaaggggaaagctggcaagattttttacgtaataacgctaaaagttttagatgtgctttactaagtcatcgcgatggagcaaaagtacatttaggtacacggcctacagaaaaacagtatgaaactctcgaaaatcaattagcctttttatgccaacaaggttttcactagagaatgcattatatgcactcagcgctgtggggcattttactttaggttgcgtattggaagatcaagagcatcaagtcgctaaagaagaaagggaaacacctactactgatagtatgccgccattattacgacaagctatcgaattatttgatcaccaaggtgcagagccagccttcttattcggccttgaattgatcatttgcggattagaaaaacaacttaaatgtgaaagtgggtcctaa |
| *luxR* | atgaaaaacataaatgccgacgacacatacagaataattaataaaattaaagcttgtagaagcaataatgatattaatcaatgcttatctgatatgactaaaatggtacattgtgaatattatttactcgcgatcatttatcctcattctatggttaaatctgatatttcaatcctagataattaccctaaaaaatggaggcaatattatgatgacgctaatttaataaaatatgatcctatagtagattattctaactccaatcattcaccaattaattggaatatatttgaaaacaatgctgtaaataaaaaatctccaaatgtaattaaagaagcgaaaacatcaggtcttatcactgggtttagtttccctattcatacggctaacaatggcttcggaatgcttagttttgcacattcagaaaaagacaactatatagatagtttatttttacatgcgtgtatgaacataccattaattgttccttctctagttgataattatcgaaaaataaatatagcaaataataaatcaaacaacgatttaaccaaaagagaaaaagaatgtttagcgtgggcatgcgaaggaaaaagctcttgggatatttcaaaaatattaggttgcagtgagcgtactgtcactttccatttaaccaatgcgcaaatgaaactcaatacaacaaaccgctgccaaagtatttctaaagcaattttaacaggagcaattgattgcccatactttaaaaattaa |
| *lacI* | gacaccatcgaatggcgcaaaacctttcgcggtatggcatgatagcgcccggaagagagtcaattcagggtggtgaatgtgaaaccagtaacgttatacgatgtcgcagagtatgccggtgtctcttatcagaccgtttcccgcgtggtgaaccaggccagccacgtttctgcgaaaacgcgggaaaaagtggaagcggcgatggcggagctgaattacattcccaaccgcgtggcacaacaactggcgggcaaacagtcgttgctgattggcgttgccacctccagtctggccctgcacgcgccgtcgcaaattgtcgcggcgattaaatctcgcgccgatcaactgggtgccagcgtggtggtgtcgatggtagaacgaagcggcgtcgaagcctgtaaagcggcggtgcacaatcttctcgcgcaacgcgtcagtgggctgatcattaactatccgctggatgaccaggatgccattgctgtggaagctgcctgcactaatgttccggcgttatttcttgatgtctctgaccagacacccatcaacagtattattttctcccatgaagacggtacgcgactgggcgtggagcatctggtcgcattgggtcaccagcaaatcgcgctgttagcgggcccattaagttctgtctcggcgcgtctgcgtctggctggctggcataaatatctcactcgcaatcaaattcagccgatagcggaacgggaaggcgactggagtgccatgtccggttttcaacaaaccatgcaaatgctgaatgagggcatcgttcccactgcgatgctggttgccaacgatcagatggcgctgggcgcaatgcgcgccattaccgagtccgggctgcgcgttggtgcggatatctcggtagtgggatacgacgataccgaagacagctcatgttatatcccgccgttaaccaccatcaaacaggattttcgcctgctggggcaaaccagcgtggaccgcttgctgcaactctctcagggccaggcggtgaagggcaatcagctgttgcccgtctcactggtgaaaagaaaaaccaccctggcgcccaatacgcaaaccgcctctccccgcgcgttggccgattcattaatgcagctggcacgacaggtttcccgactggaaagcgggcagtga |
| *gfp* | atgcgtaaaggagaagaacttttcactggagttgtcccaattcttgttgaattagatggtgatgttaatgggcacaaattttctgtcagtggagagggtgaaggtgatgcaacatacggaaaacttacccttaaatttatttgcactactggaaaactacctgttccatggccaacacttgtcactactttcggttatggtgttcaatgctttgcgagatacccagatcatatgaaacagcatgactttttcaagagtgccatgcccgaaggttatgtacaggaaagaactatatttttcaaagatgacgggaactacaagacacgtgctgaagtcaagtttgaaggtgatacccttgttaatagaatcgagttaaaaggtattgattttaaagaagatggaaacattcttggacacaaattggaatacaactataactcacacaatgtatacatcatggcagacaaacaaaagaatggaatcaaagttaacttcaaaattagacacaacattgaagatggaagcgttcaactagcagaccattatcaacaaaatactccaattggcgatggccctgtccttttaccagacaaccattacctgtccacacaatctgccctttcgaaagatcccaacgaaaagagagaccacatggtccttcttgagtttgtaacagctgctgggattacacatggcatggatgaactatacaaataa |
| *gfp*$_{ASV}$ | atgcgtaaaggagaagaacttttcactggagttgtcccaattcttgttgaattagatggtgatgttaatgggcacaaattttctgtcagtggagagggtgaaggtgatgcaacatacggaaaacttacccttaaatttatttgcactactggaaaactacctgttccatggccaacacttgtcactactttcggttatggtgttcaatgctttgcgagatacccagatcatatgaaacagcatgactttttcaagagtgccatgcccgaaggttatgtacaggaaagaactatatttttcaaagatgacgggaactacaagacacgtgctgaagtcaagtttgaaggtgatacccttgttaatagaatcgagttaaaaggtattgattttaaagaagatggaaacattcttggacacaaattggaatacaactataactcacacaatgtatacatcatggcagacaaacaaaagaatggaatcaaagttaacttcaaaattagacacaacattgaagatggaagcgttcaactagcagaccattatcaacaaaatactccaattggcgatggccctgtccttttaccagacaaccattacctgtccacacaatctgccctttcgaaagatcccaacgaaaagagagaccacatggtccttcttgagtttgtaacagctgctgggattacacatggcatggatgaactatacaaaaggcctgcagcaaacgacgaaaactacgctgcatcagtttaa |
| *mCherry* | atggtgagcaagggcgaggaggataacatggccatcatcaaggagttcatgcgcttcaaggttcacatggagggctccgtgaacggccacgagttcgagatcgagggcgagggcgagggccgcccctacgagggcacccagaccgccaagctgaaggtgaccaagggtggcccccctgcccttcgcctgggacatcctgtcccctcagttcatgtacggctccaaggcctacgtgaagcaccccgccgacatccccgactacttgaagctgtccttccccgagggcttcaagtgggagcgcgtgatgaacttcgaggacggcggcgtggtgaccgtgacccaggactcctccctgcaagacggcgagttcatctacaaggtgaagctgcgcggcaccaacttcccctccgacggccccgtaatgcagaagaagactatgggctgggaggcctcctccgagcggatgtaccccgaggacggcgcgctgaagggcgagatcaagcagaggctgaagctgaaggacggcggccactacgacgctgaggtcaagaccacctacaaggccaagaagcccgtgcaactgcccggcgcgtacaacgtcaacatcaagttggacatcacctcccacaacgaggactacaccatcgtggaacagtacgaacgcgccgagggccgccactccaccggcggcatggacgagctgtataagtaa |

| | |
|---|---|
| *mCherry-RT* | atggtgagcaagggcgaggaggataacatggccatcatcaaggagttcatgcgcttcaaggttcacatggagggctccgtgaacggcca<br>cgagttcgagatcgagggcgagggcgagggccgcccctacgagggcacccagaccgccaagctgaaggtgaccaagggtggcccc<br>ctgcccttcgcctgggacatcctgtcccctcagttcatgtacggctccaaggcctacgtgaagcaccccgccgacatccccgactacttga<br>agctgtccttccccgagggcttcaagtgggagcgcgtgatgaacttcgaggacggcggcgtggtgaccgtgacccaggactcctccctg<br>caagacggcgagttcatctacaaggtgaagctgcgcggcaccaacttcccctccgacggccccgtaatgcagaagaagactatgggct<br>gggaggcctcctccgagcggatgtaccccgaggacggcgcgctgaagggcgagatcaagcagaggctgaagctgaaggacggcgg<br>ccactacgacgctgaggtcaagaccacctacaaggccaagaagcccgtgcaactgcccggcgcgtacaacgtcaacatcaagttggac<br>atcacctcccacaacgaggactacaccatcgtggaacagtacgaacgcgccgagggccgccactccaccggcggcatggacgagctg<br>tataagggcggtggcggttctggcggtggcggtagcaaatccgctgaatatttgaacacttttagattgagaaatctcggcctacctgtcatg<br>aacaatttgcatgacatgtctaaggcgactcgcatatctgttgaaacacttcggttgttaatctatacagctgattttcgctataggatctacact<br>gtagaaaagaaaggcccagagaagagaatgagaaccatttaccaaccttctcgagaacttaaagccttacaaggatgggttctacgtaac<br>attttagataaactgtcgtcatctcctttttctattggatttgaaaagcaccaatctattttgaataatgctaccccgcatattggggcaaactttat<br>actgaatattgatttggaggatttttttcccaagtttaactgctaacaaagtttttggagtgttccattctcttggttataatcgactaatatcttcagtt<br>ttgacaaaaatatgttgttataaaaatctgctaccacaaggtgctccatcatcacctaaattagctaatctaatatgttctaaacttgattatcgtat<br>tcagggttatgcaggtagtcggggcttgatatatacgagatatgccgatgatctcaccttatctgcacagtctatgaaaaaggttgttaaagc<br>acgtgatttttttattttctataatcccaagtgaaggattggttattaactcaaaaaaaacttgtattagtgggcctcgtagtcagaggaaagttac<br>aggtttagttatttcacaagagaaagttgggataggtagagaaaaatataaagaaattagagcaaagatacatcatatattttgcggtaagtct<br>tctgagatagaacacgttaggggatggttgtcatttattttaagtgtggattcaaaaagccataggagattaataacttatattagcaaattaga<br>aaaaaaatatggaaagaaccctttaaataaagcgaagacctaa |
| *RT-mCherry* | atgaaatccgctgaatatttgaacacttttagattgagaaatctcggcctacctgtcatgaacaatttgcatgacatgtctaaggcgactcgcat<br>atctgttgaaacacttcggttgttaatctatacagctgattttcgctataggatctacactgtagaaaagaaaggcccagagaagagaatgag<br>aaccatttaccaaccttctcgagaacttaaagccttacaaggatgggttctacgtaacattttagataaactgtcgtcatctcctttttctattgga<br>tttgaaaagcaccaatctattttgaataatgctaccccgcatattggggcaaactttatactgaatattgatttggaggatttttttcccaagtttaa<br>ctgctaacaaagtttttggagtgttccattctcttggttataatcgactaatatcttcagttttgacaaaaatatgttgttataaaaatctgctaccac<br>aaggtgctccatcatcacctaaattagctaatctaatatgttctaaacttgattatcgtattcagggttatgcaggtagtcggggcttgatatata<br>cgagatatgccgatgatctcaccttatctgcacagtctatgaaaaaggttgttaaagcacgtgatttttttattttctataatcccaagtgaaggat<br>tggttattaactcaaaaaaaacttgtattagtgggcctcgtagtcagaggaaagttacaggtttagttatttcacaagagaaagttgggatagg<br>tagagaaaaatataaagaaattagagcaaagatacatcatatattttgcggtaagtcttctgagatagaacacgttaggggatggttgtcattt<br>attttaagtgtggattcaaaaagccataggagattaataacttatattagcaaattagaaaaaaaatatggaaagaaccctttaaataaagcga<br>agaccggcggtggcggttctggcggtggcggtagcgtgagcaagggcgaggaggataacatggccatcatcaaggagttcatgcgctt<br>caaggttcacatggagggctccgtgaacggccacgagttcgagatcgagggcgagggcgagggccgcccctacgagggcacccaga<br>ccgccaagctgaaggtgaccaagggtggcccccctgcccttcgcctgggacatcctgtcccctcagttcatgtacggctccaaggcctacg<br>tgaagcaccccgccgacatccccgactacttgaagctgtccttccccgagggcttcaagtgggagcgcgtgatgaacttcgaggacggc<br>ggcgtggtgaccgtgacccaggactcctccctgcaagacggcgagttcatctacaaggtgaagctgcgcggcaccaacttcccctccga<br>cggccccgtaatgcagaagaagactatgggctgggaggcctcctccgagcggatgtaccccgaggacggcgcgctgaagggcgaga<br>tcaagcagaggctgaagctgaaggacggcggccactacgacgctgaggtcaagaccacctacaaggccaagaagcccgtgcaactg<br>cccggcgcgtacaacgtcaacatcaagttggacatcacctcccacaacgaggactacaccatcgtggaacagtacgaacgcgccgagg<br>gccgccactccaccggcggcatggacgagctgtataag |
| *tar-tetR-gfp* | atgattaaccgtatccgcgtagtcacgctgttggtaatggtgctgggggtattcgcactgttacagcttatttccggcagtctgtttttttcttccc<br>ttcaccatagccagaagagctttgtggtttccaatcaattacgggaacagcagggcgagctgacgtcaacctgggatttaatgctgcaaac<br>gcgcattaacctgagtcgttcagcggtacggatgatgatggattcctccaatcaacaaagtaacgccaaagttgaattgctcgatagcgcc<br>aggaaaacattggcgcaggcagcgacgcattataaaaaattcaaaagcatggcaccgttacctgaaatggtcgctaccagtcgtaatattg<br>atgaaaaatataaaaactattacacagcgttaactgaactgattgattatctagattatggcaatactggagcttatttcgctcagccaacccag<br>ggaatgcaaaatgcaatgggcgaagcgtttgctcagtacgccctcagcagtgaaaaactgtatcgcgatatcgtcactgacaacgcagat<br>gattaccgatttgcccagtggcaactggcggttatcgcgctggtggtggtattgattctgctggtggcgtggtacggcattcgccgtatgttg<br>cttactccgctggcaaaaattattgctcacattcgcgaaatcgccggtggtaacctggcgaataccctgaccattgacgggcgcagtgaaa<br>tgggcgacctggcgcagagcgtttcacatatgcaacgctctttgactgacaccgtcactcatgtccgcgaaggttcagatgccatctatgcc<br>ggtacccgtgaaattgcggcgggcaacaccgatctttcctcccgtactgaacagcaggcatccgcgctggaagaaactgccgccagcat<br>ggagcagctcaccgcgacagtgaagcaaaacgccgataacgcccgccaggcctcgcaactggcgcaaagtgcctccgacaccgccc<br>agcacggcggcaaagtggtggatggcgtagtgaaaacgatgcatgagatcgccgatagttcgaagaaaattgccgacattatcagcgtta<br>tcgacggtattgccttccagactaatatcctcgcgctgaatgccgcggttgaagccgcgcgtgcgggtgaacagggccgtggttttgccgt<br>ggtggcgggtgaagtgcgtaatcttgccagtcgcagcgcccaggcggcaaaagagatcaaagccctcattgaagactccgtctcacgc<br>gttgataccggttcggtgctggtcgaaagcgccggggaaacaatgaacaatatcgtcaatgctgtcactcgcgtgactgacattatgggcg<br>agattgcatcggcatcggatgaacagagccgtggcatcgatcaagtcgcattggcggtttcggaaatggatcgcgtcacgcaacagaac<br>gcatcgctggtgcaggaatcagctgccgccgccgctgcgctggaagaacaggcgagtcgtttaacgcaagcagtttccgcgttccgtct<br>ggcagccagcccactcaccaataaaccgcaaacaccatcccgtcctgccagtgagcaaccaccggctcagccacgactgcgaattgct<br>gaacaagatccaaactgggaaacatttggcggtggcggttctggcggtggcggtagctccagattagataaaagtaaagtgattaacagc |

| | |
|---|---|
| | gcattagagctgcttaatgaggtcggaatcgaaggtttaacaacccgtaaactcgcccagaagctaggtgtagagcagcctacattgtattg<br>gcatgtaaaaaataagcgggctttgctcgacgccttagccattgagatgttagataggcaccatactcacttttgccctttagaaggggaaa<br>gctggcaagatttttacgtaataacgctaaaagttttagatgtgctttactaagtcatcgcgatggagcaaaagtacatttaggtacacggcct<br>acagaaaaacagtatgaaactctcgaaaatcaattagcctttttatgccaacaaggtttttcactagagaatgcattatatgcactcagcgctgt<br>ggggcattttactttaggttgcgtattggaagatcaagagcatcaagtcgctaaagaagaaagggaaacacctactactgatagtatgccgc<br>cattattacgacaagctatcgaattatttgatcaccaaggtgcagagccagccttcttattcggccttgaattgatcatttgcggattagaaaaa<br>caacttaaatgtgaaagtgggtcctccatggtgagcaaaggagaagaacttttcactggagttgtcccaattcttgttgaattagatggtgatg<br>ttaatgggcacaaattttctgtcagtggagagggtgaaggtgatgcaacatacggaaaacttacccttaaatttatttgcactactggaaaact<br>acctgttccatggccaacacttgtcactactttcggttatggtgttcaatgctttgcgagatacccagatcatatgaaacagcatgactttttcaa<br>gagtgccatgcccgaaggttatgtacaggaaagaactatatttttcaaagatgacgggaactacaagacacgtgctgaagtcaagtttgaa<br>ggtgatacccttgttaatagaatcgagttaaaaggtattgattttaaagaagatggaaacattcttggacacaaattggaatacaactataactc<br>acacaatgtatacatcatggcagacaaacaaaagaatggaatcaaagttaacttcaaaattagacacaacattgaagatggaagcgttcaa<br>ctagcagaccattatcaacaaaatactccaattggcgatggccctgtccttttaccagacaaccattacctgtccacacaatctgccctttcga<br>aagatcccaacgaaaagagagaccacatggtccttcttgagtttgtaacagctgctgggattacacatggcatggatgaactatacaaataa |
| *tsr-tetR-gfp* | atgttaaaacgtatcaaaattgtgaccagcttactgctggttttggccgtttttggccttttacaactgacatcaggcggtctgttctttaatgcctt<br>aaagaatgacaaagaaaatttcactgttttacaaaccattcgccagcagcaatccacgctgaatggcagctgggtcgcgttgttgcagacg<br>cgtaacaccctcaaccgcgcgggtatccgctacatgatggatcagaataatattggtagcggttcaaccgttgctgagctgatggagagtg<br>ccagtatttcgctgaaacaggcggaaaaaaactgggcggattacgaagcgttgccgcgtgacccgcgtcagagcaccgccgcagcgg<br>cagagatcaaacgtaattacgatatttatcacaatgcgctggcggagctgatccaactgttaggtgcaggcaaaatcaacgagttctttgatc<br>agccgacccagggatatcaggacggtttcgagaagcagtatgtggcttacatggagcaaaacgatcggctccatgatatcgccgtcagc<br>gataacaatgcctcctacagccaggcgatgtggattctggtgggcgtgatgatcgtcgtactggcggtcatcttcgccgtctggttcggtatt<br>aaagcctcgctggtagcgccaatgaatcgcctgattgacagcattcgtcatattgcaggcggcgatctggtgaaaccgattgaggtggatg<br>gctctaatgagatggggcaactggcagagagtttgcgccatatgcagggagagctgatgcgtaccgtcggtgatgtgcgcaacggggc<br>caatgccatctatagcggtgccagcgaaatcgccaccggcaataacgatctctcttcgcgcaccgagcaacaggccgcttcgctggaag<br>agacggcagccagcatggagcaactgaccgcaacggtgaagcagaacgccgaaaatgcgcgccaggccagccatctggcgttaagt<br>gcttctgaaacggcgcaacgcggcggtaaagtggtagataacgtggtgcagactatgcgcgatatctccaccagttcgcagaaaatcgc<br>cgatattatcagcgtaattgacggcattgccttccagaccaatattctggctttgaacgcggcggttgaggctgcgcgtgcgggtgagcaa<br>gggcgcggttttgcggtggtcgcgggagaagtgcgtaatctggcccagcgcagcgcccaggcggctcgtgaaattaaaagcctgattg<br>aagactcggtggggaaagtggatgttggctctacgctggtcgaaagcgccggggaaacaatggcggagattgtcagcgccgtgacccg<br>cgtgacggacattatgggcgaaattgcttctgcttctgatgagcagagccgtggtatcgatcaggttggcttagcggttgctgagatggacc<br>gggtaactcaacagaacgccgcgctggtggaagagtctgccgctgccgccgccgcgctggaagagcaggccagtcgcctgaccgaa<br>gcagtggcagtgttccggattcagcaacagcagcgtgaaacatcggctgtggtaaaaaccgtgacgccagctgcgccgcgtaaaatgg<br>ccgtggcagatagcgaggagaactgggaaacatttggcggtggcggttctggcggtggcggtagctccagattagataaaagtaaagtg<br>attaacagcgcattagagctgcttaatgaggtcggaatcgaaggtttaacaacccgtaaactcgcccagaagctaggtgtagagcagccta<br>cattgtattggcatgtaaaaaataagcgggctttgctcgacgccttagccattgagatgttagataggcaccatactcacttttgccctttagaa<br>ggggaaagctggcaagatttttacgtaataacgctaaaagttttagatgtgctttactaagtcatcgcgatggagcaaaagtacatttaggta<br>cacggcctacagaaaaacagtatgaaactctcgaaaatcaattagcctttttatgccaacaaggtttttcactagagaatgcattatatgcact<br>cagcgctgtggggcattttactttaggttgcgtattggaagatcaagagcatcaagtcgctaaagaagaaagggaaacacctactactgata<br>gtatgccgccattattacgacaagctatcgaattatttgatcaccaaggtgcagagccagccttcttattcggccttgaattgatcatttgcgga<br>ttagaaaaacaacttaaatgtgaaagtgggtcctccatggtgagcaaaggagaagaacttttcactggagttgtcccaattcttgttgaattag<br>atggtgatgttaatgggcacaaattttctgtcagtggagagggtgaaggtgatgcaacatacggaaaacttacccttaaatttatttgcactac<br>tggaaaactacctgttccatggccaacacttgtcactactttcggttatggtgttcaatgctttgcgagatacccagatcatatgaaacagcat<br>gactttttcaagagtgccatgcccgaaggttatgtacaggaaagaactatatttttcaaagatgacgggaactacaagacacgtgctgaagt<br>caagtttgaaggtgatacccttgttaatagaatcgagttaaaaggtattgattttaaagaagatggaaacattcttggacacaaattggaatac<br>aactataactcacacaatgtatacatcatggcagacaaacaaaagaatggaatcaaagttaacttcaaaattagacacaacattgaagatgg<br>aagcgttcaactagcagaccattatcaacaaaatactccaattggcgatggccctgtccttttaccagacaaccattacctgtccacacaatc<br>tgccctttcgaaagatcccaacgaaaagagagaccacatggtccttcttgagtttgtaacagctgctgggattacacatggcatggatgaac<br>tatacaaataa |
| *tatA-tetR-gfp* | atgggtggtatcagtatttggcagttattgattattgccgtcatcgttgtactgctttttggcaccaaaaagctcggctccatcggttccgatctt<br>ggtgcgtcgatcaaaggctttaaaaaagcaatgagcgatgatgaaccaaagcaggataaaaccagtcaggatgctgattttactgcgaaa<br>actatcgccgataagcaggcggatacgaatcaggaacaggctaaaacagaagacgcgaagcgccacgataaagagcaggtgggcgg<br>tggcggttctggcggtggcggtagctccagattagataaaagtaaagtgattaacagcgcattagagctgcttaatgaggtcggaatcgaa<br>ggtttaacaacccgtaaactcgcccagaagctaggtgtagagcagcctacattgtattggcatgtaaaaaataagcgggctttgctcgacg<br>ccttagccattgagatgttagataggcaccatactcacttttgccctttagaaggggaaagctggcaagatttttacgtaataacgctaaaag<br>ttttagatgtgctttactaagtcatcgcgatggagcaaaagtacatttaggtacacggcctacagaaaaacagtatgaaactctcgaaaatca<br>attagcctttttatgccaacaaggtttttcactagagaatgcattatatgcactcagcgctgtggggcattttactttaggttgcgtattggaagat<br>caagagcatcaagtcgctaaagaagaaagggaaacacctactactgatagtatgccgccattattacgacaagctatcgaattatttgatca |

| | |
|---|---|
| | ccaaggtgcagagccagccttcttattcggccttgaattgatcatttgcggattagaaaaacaacttaaatgtgaaagtgggtcctccatggtgagcaaaggagaagaacttttcactggagttgtcccaattcttgttgaattagatggtgatgttaatgggcacaaattttctgtcagtggagagggtgaaggtgatgcaacatacggaaaacttacccttaaatttatttgcactactggaaaactacctgttccatggccaacacttgtcactactttcggttatggtgttcaatgctttgcgagatacccagatcatatgaaacagcatgactttttcaagagtgccatgcccgaaggttatgtacaggaaagaactatatttttcaaagatgacgggaactacaagacacgtgctgaagtcaagtttgaaggtgatacccttgttaatagaatcgagttaaaaggtattgattttaaagaagatggaaacattcttggacacaaattggaatacaactataactcacacaatgtatacatcatggcagacaaacaaaagaatggaatcaaagttaacttcaaaattagacacaacattgaagatggaagcgttcaactagcagaccattatcaacaaaatactccaattggcgatggccctgtccttttaccagacaaccattacctgtccacacaatctgccctttcgaaagatcccaacgaaaagagagaccacatggtccttcttgagtttgtaacagctgctgggattacacatggcatggatgaactatacaaataa |
| *popZ-tetR-gfp* | atgtccgatcagtctcaagaacctacaatggaggaaatcctcgcctccattcgacgcatcatctcggaggatgacgcgccggcggagcctgcggccgaagcggcgcccccgccgccgccggaacccgaacctgaaccggtgtcgttcgacgacgaggttctggaattgacggacccgatcgcgcccgagcccgagctgccgccgctggagactgtcggcgacatcgacgtctattcgccgccggaacctgagtcggaaccggcctacacgccgccgccggcggctccggtgtttgatcgcgacgaagtcgccgagcagctggtcggcgtttcggccgcttcggccgcggcgagcgccttcggcagcctgagctcggccctgctgatgcccaaggacggtcggacgctggaagacgtcgtacgcgagctgctgcgcccgctgctcaaggagtggctggaccagaacctgccgcgcatcgtcgagaccaaggttgaggaagaagtgcagcgtatctctcggggacgcggcgccggcggtggcggttctggcggtggcggtagctccagattagataaaagtaaagtgattaacagcgcattagagctgcttaatgaggtcggaatcgaaggtttaacaacccgtaaactcgcccagaagctaggtgtagagcagcctacattgtattggcatgtaaaaaataagcgggctttgctcgacgccttagccattgagatgttagataggcaccatactcacttttgccctttagaaggggaaagctggcaagatttttacgtaataacgctaaaagttttagatgtgctttactaagtcatcgcgatggagcaaaagtacatttaggtacacggcctacagaaaaacagtatgaaactctcgaaaatcaattagcctttttatgccaacaaggtttttcactagagaatgcattatatgcactcagcgctgtggggcattttactttaggttgcgtattggaagatcaagagcatcaagtcgctaaagaagaaagggaaacacctactactgatagtatgccgccattattacgacaagctatcgaattatttgatcaccaaggtgcagagccagccttcttattcggccttgaattgatcatttgcggattagaaaaacaacttaaatgtgaaagtgggtcctccatggtgagcaaaggagaagaacttttcactggagttgtcccaattcttgttgaattagatggtgatgttaatgggcacaaattttctgtcagtggagagggtgaaggtgatgcaacatacggaaaacttacccttaaatttatttgcactactggaaaactacctgttccatggccaacacttgtcactactttcggttatggtgttcaatgctttgcgagatacccagatcatatgaaacagcatgactttttcaagagtgccatgcccgaaggttatgtacaggaaagaactatatttttcaaagatgacgggaactacaagacacgtgctgaagtcaagtttgaaggtgatacccttgttaatagaatcgagttaaaaggtattgattttaaagaagatggaaacattcttggacacaaattggaatacaactataactcacacaatgtatacatcatggcagacaaacaaaagaatggaatcaaagttaacttcaaaattagacacaacattgaagatggaagcgttcaactagcagaccattatcaacaaaatactccaattggcgatggccctgtccttttaccagacaaccattacctgtccacacaatctgccctttcgaaagatcccaacgaaaagagagaccacatggtccttcttgagtttgtaacagctgctgggattacacatggcatggatgaactatacaaataa |

### Table S4. DNA oligos used in this study.

| **Name** | **Sequence** |
|---|---|
| Seq_1 | ataacgcggcactcatgattatcatcactcttttagtccctattacagatcggcatcctgcattgaatctgagttactgtctg |
| Seq_2 | tccgccgcaagatctagtgcaatgggtcagaaaaaacggg |
| Seq_3 | gtaagaccgctctgttccgacaccaccataacgcggcactc |
| qPCR_plasmid_F | caacctctggatgttgtttcg |
| qPCR_plasmd_R | gcggatttcatgaaagttgc |
| qPCR_rtDNA_F | tctgagttactgtctgttttcct |
| qPCR_rtDNA_R | gtcagaaaaaacgggtttcct |
| WT retron-DNA oligo | gtcagaaaaaacgggtttcctggttggctcggagagcatcaggcgatgctctccgttccaacaaggaaaacagacagtaactcaga |
